# Phylogenetic diversity of two common *Trypanosoma cruzi* lineages in the Southwestern United States

**DOI:** 10.1101/2021.11.10.468115

**Authors:** Carlos A. Flores-López, Elizabeth A. Mitchell, Carolina E. Reisenman, Sahotra Sarkar, Philip C. Williamson, Carlos A. Machado

**Affiliations:** Department of Biology, University of Maryland, College Park, Maryland, USA; Facultad de Ciencias, Universidad Autónoma de Baja California, Ensenada, Baja California, Mexico; Tick-Borne Disease Research Laboratory, Department of Microbiology, Immunology, and Genetics, University of North Texas Health Science Center, Fort Worth, Texas, USA; Department of Pathology and Microbiology, University of Nebraska Medical Center, Omaha, Nebraska, USA; Department of Molecular and Cell Biology, University of California, Berkeley, California, USA; Department of Philosophy and Department of Integrative Biology, University of Texas, Austin, Texas, USA; Creative Testing Solutions, Tempe, Arizona, USA

**Keywords:** *Trypanosoma cruzi*, Chagas Disease, DTU, genetic diversity, evolution, triatomines

## Abstract

*Trypanosoma cruzi* is the causative agent of Chagas disease, a devastating parasitic disease endemic to Central and South America, Mexico, and the USA. We characterized the genetic diversity of *T. cruzi* circulating in five triatomine species (*Triatoma gerstaeckeri, T. lecticularia, T. indictiva, T. sanguisuga* and *T. recurva*) collected in Texas and Southern Arizona using nucleotide sequences from four single-copy loci (COII-ND1, MSH2, DHFR-TS, TcCLB.506529.310). All *T. cruzi* variants fall in two main genetic lineages: 75% of the samples corresponded to *T. cruzi* Discrete Typing Unit (DTU) I (TcI), and 25% to a North American specific lineage previously labelled TcIV-USA. Phylogenetic and sequence divergence analyses of our new data plus all previously published sequence data from those 4 genes collected in the USA, show that TcIV-USA is significantly different from any other previously defined *T. cruzi* DTUs. The significant level of genetic divergence between TcIV-USA and other *T. cruzi* lineages should lead to an increased focus on understanding the epidemiological importance of this lineage, as well as its geographical range and pathogenicity in humans and domestic animals. Our findings further corroborate the fact that there is a high genetic diversity of the parasite in North America and emphasize the need for appropriate surveillance and vector control programs for Chagas disease in southern USA and Mexico.

## 1. INTRODUCTION

Chagas disease is caused by the protozoan parasite *Trypanosoma cruzi*, which is transmitted to humans by blood-sucking insects of the family Reduviidae (Triatominae). This is the most important parasitic disease in the Americas, with about 6-7 million people infected and 75 million people considered at risk, mostly in Mexico, Central and South America (WHO, 2019). Between 10-30% percent of infected people eventually develop illness of the cardiac, gastrointestinal and/or nervous systems, resulting in severe debilitation, economic burden, and ultimately death (Barrett et al., 2003; Prata, 2001). Chagas is mainly a vector-borne disease in which the parasite is transmitted when feces and/or urine from infected insects contact the skin or oral and/or eye mucous membranes.

It is recognized that Chagas disease, traditionally confined to Mexico, Central and South America, is becoming an important health issue not only in the USA but also in other countries worldwide (Coura and Vinas, 2010; Lidani et al., 2019; Manne-Goehler et al., 2016; Montgomery et al., 2014). In the USA, it is estimated that between 250,000 and 300,000 people, mostly individuals born in countries where the disease is endemic, are infected with *T. cruzi* (Bern et al., 2019; Bern and Montgomery, 2009; Manne-Goehler et al., 2016; Montgomery et al., 2014; Yadon and Schmunis, 2009). These numbers are significant and emphasize the need for recognition among physicians and public health officials, as the parasite can be transmitted through blood transfusion and congenitally (Manne-Goehler et al., 2015; Parise et al., 2014; Verani et al., 2010). In addition, both infected triatomines and mammalian reservoirs of the parasite such as packrats, armadillos, raccoons, skunks, foxes, mice, opossums, dogs and captive nonhuman primates are plentiful in the USA, particularly in southern states (Aleman et al., 2017; Barnabe et al., 2001; Beard et al., 2003; Bern et al., 2011; Bern et al., 2019; Bradley et al., 2000; Curtis-Robles et al., 2018; Elmayan et al., 2019; Ghersi et al., 2020; Herrera et al., 2019; Hodo et al., 2018; Kjos et al., 2008; Meurs et al., 1998; Padilla et al., 2021; Pung et al., 1995; Rodriguez et al., 2021; Roellig et al., 2008; Shender et al., 2016; Vandermark et al., 2018; Zecca et al., 2020). Despite this, only a small (< 100), but increasing number of autochthonous vector-borne cases of *T. cruzi* transmission have been reported to date, all in the southern part of the country (Reviewed in (Bern et al., 2019; Lynn et al., 2020). Interestingly, some individual blood donors that tested positive for the presence of the parasite never left the USA (Cantey et al., 2012), indicating that autochthonous transmission may be even higher and undiscovered. In addition, many Chagas disease cases might have been overlooked because the early phase of the infection is often asymptomatic, and because some chagasic cardiomyopathies may be misdiagnosed (Herwaldt et al., 2000; Kirchhoff, 1993; Milei et al., 2009).

Transmission of *T. cruzi* in the USA mostly occurs within sylvatic cycles (Bern et al., 2019) although triatomines, often infected, occasionally invade human dwellings during dispersing flights and reportedly become in contact with humans and pets (Beard et al., 2003; Klotz et al., 2010; Lynch and Pinnas, 1978; Moffitt et al., 2003; Reisenman et al., 2010; Reisenman et al., 2012; Stevens et al., 2012). Although triatomines in the USA appear to have a relatively low capacity for domiciliation and human transmission (Klotz et al., 2016), domestic cycles of disease transmission involving infected insects, dogs and sylvatic reservoirs have been reported in southern Texas (Beard et al., 2003; Garcia et al., 2016). In particular, the finding that dogs in the USA have increasing roles in enzootic cycles (Bradley et al., 2000; Curtis-Robles et al., 2017; Elmayan et al., 2019; Garcia et al., 2016; Kjos et al., 2008), along with the fact these animals are important domestic reservoirs of *T. cruzi* (Crisantes et al., 2006; Curtis-Robles et al., 2017; Gürtler et al., 2007; Gurtler et al., 1991), is highly relevant. The presence of infected triatomines, small mammals such as packrats, and dogs that spend time outside the house establishes a scenario in which domestic transmission cycles of the parasite are possible.

Currently, the parasite’s genetic diversity is divided into seven lineages or Discrete Typing Units (DTUs) termed TcI-TcVI and TcBat (Brisse et al., 2000; Pinto et al., 2012; Zingales et al., 2012). Although asexuality has been traditionally assumed to be the most common mechanism by which the parasite reproduces (Tibayrenc and Ayala, 1987, 1988), recent studies have shown that sexual reproduction happens in nature and that clonal and sexual panmictic populations can coexist (Berry et al., 2019; Schwabl et al., 2019). Furthermore, it is well-known that genetic exchange events have played important roles generating genetic diversity in this parasite. For instance, two of the main lineages with widespread geographic distribution are hybrids (TcV-TcVI), the result of an ancient hybridization event that took place between the ancestors of DTUs TcII and TcIII (Brisse et al., 2003; Flores-López and Machado, 2011; Machado and Ayala, 2001, 2002), and surveys of variation using mitochondrial and nuclear genes often show evidence of genetic exchange (Lewis et al., 2011; Messenger et al., 2012; Roellig et al., 2013; Shender et al., 2016). The findings of an additional genetic lineage that infects bats in Central America (TcBat) (Lima et al., 2015; Pinto et al., 2012), and the more complex genetic structure in TcI (Llewellyn et al., 2009; Zumaya-Estrada et al., 2012), suggest that the genetic diversity of *T. cruzi* may not have been thoroughly sampled. As sampling is expanded to more regions, a more complex view of the genetic diversity of the parasite may be revealed.

Studies attempting to uncover the genetic diversity of *T. cruzi* in the USA have become more common in recent years. TcI and TcIV are the most prevalent DTUs found in the USA (Barnabe et al., 2001; Breniere et al., 2016; Curtis-Robles et al., 2018; Curtis-Robles et al., 2017; Garcia et al., 2017; Herrera et al., 2015; Meyers et al., 2017; Roellig et al., 2008; Roellig et al., 2013; Shender et al., 2016; Vandermark et al., 2018). TcI was thought to be the only DTU involved in autochthonous human infections in this country (Roellig et al., 2008), but a recent study uncovered the presence of non-TcI DTUs and mixed TcI-nonTcI infections in Chagas disease patients from Texas which acquired the infection autochthonously (Garcia et al., 2017). Additionally, mixed TcI/TcIV infections and evidence of genetic exchange between TcI and TcIV have been detected in wildlife reservoirs (Curtis-Robles et al., 2018; Roellig et al., 2013). Furthermore, TcVI was recently detected in captive primates (Herrera et al., 2019), and TcII was also detected in rodents from Louisiana (Herrera et al., 2015). More recently, all DTUs except TcIII and TcBat were detected in a small sample of rodents from New Orleans using next-generation sequencing of the mini exon gene (Pronovost et al., 2020).

All the above discoveries beseech for a better understanding of the genetic diversity of *T. cruzi* currently present in the USA. We thus conducted a phylogenetic study of *T. cruzi* sequences isolated from triatomine bugs from two southern USA states, Arizona and Texas, where infected triatomines are readily abundant and reportedly feed on humans and pets (Beatty et al., 2018; Garcia et al., 2015; Garcia et al., 2016; Reisenman et al., 2010; Stevens et al., 2012). We used a combination of three single copy nuclear genes and one mitochondrial region that provide more resolution to identify DTUs and uncover phylogenetic relationships than data from surveys conducted with single or less variable regions. We found two genetic lineages of *T. cruzi* circulating in the sampled Triatomines, one of which has significant genetic divergence from previously defined *T. cruzi* DTUs.

## 2. MATERIALS AND METHODS

### 2.1 Collection and location of Triatomine samples

Hand collection, dry ice traps, white light, and UV traps were used for Triatomine collections across Texas and Arizona (Fig. 1; Table S1) (Mitchell, 2013; Reisenman et al., 2010). Fifty-five individual Triatomine bugs from Texas and Arizona that were infected with *T. cruzi* are the focus of this study. Collection records for those individual insects were previously reported (Mitchell, 2013; Reisenman et al., 2012) but detailed location information is presented in S1 Table. In Texas, the majority of infected specimens were *T. gerstaeckeri* (n=44), with a lower frequency of *T. lecticularia* (n=4), *T. sanguisuga* (n=2) and *T. indictiva* (n=1) (Mitchell, 2013). From Arizona we only collected data from infected *T. recurva* (n=4) (Reisenman et al., 2010; Reisenman et al., 2012). Collected insects were individually placed in 95% ethanol immediately after collection or upon death and stored at 4°C until analysis. The posterior end of each *Triatoma* specimen was removed to isolate the hindgut. DNA was extracted using the QIAGEN Blood and Tissue kit following the manufacturer’s protocol.

**Fig. 1.**
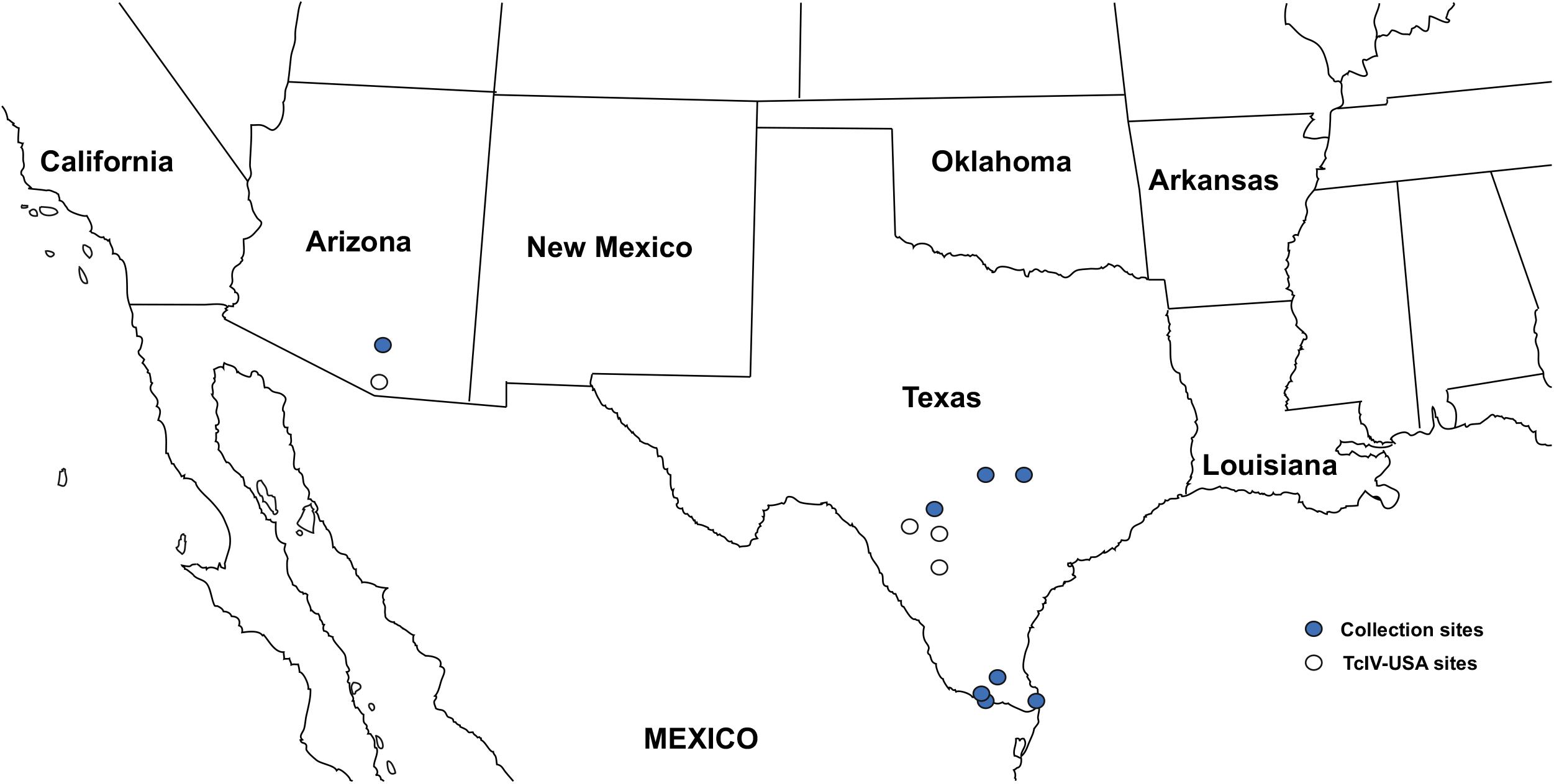
Southern USA map with collection sites for Triatomines used in this study. Collection sites for specimens of Triatomine vectors (*Triatoma gerstaeckeri, T. lecticularia, T.indictiva, T. sanguisuga* and *T. recurva)* collected in southern Arizona and Texas. Geographic coordinates of each site can be found in Table S1.

### 2.2 PCR amplification and sequencing

We conducted Multilocus sequence typing (MLST) of all the Triatomine DNA samples using previously published primers that target four variable *T. cruzi* single-copy genes: a mitochondrial region (1,226 bp) encompassing the maxicircle-encoded genes cytochrome oxidase subunit II and NADH dehydrogenase subunit 1 (COII-ND1) (Machado and Ayala, 2001); a partial segment (727 bp) of nuclear gene TcCLB.506529.310 (Former ID: Tc00.1047053506529.310) part of the Ydr279p protein family (RNase H2 complex component) (Flores-López and Machado, 2011); a 737 bp region of the mismatch repair gene (MSH2) (Augusto-Pinto et al., 2001); and a 1,408 bp region of the Dihydrofolate reductase-thymidylate synthase (DHFR-TS) gene (Machado and Ayala, 2002). The rationale for selecting these four single-copy genes was based on their previous use as taxonomic markers for *T. cruzi* and the availability of reference sequences of each DTU for the level of resolution needed in this study. Conditions for the PCR amplifications were: 35 cycles of a 30 second denaturation step at 94°C, annealing at 55°C and 58°C for 30 seconds for the nuclear and mitochondrial loci respectively, and extension at 68°C for 60 and 90 seconds respectively. Amplicons were prepared for sequencing as previously described (Mitchell, 2013), and PCR primers were used for bidirectional sequencing on a 3130*xl* Genetic Analyzer (Applied Biosystems). Sequences were edited using CodonCode Aligner (CodonCode Corporation) and deposited in GenBank (Accession Nos. COII-NDI: MF670300-MF670349, DHFR-TS: MF784866-MF784877, MSH2: MF074143-MF074183, TcCLB.506529.310: MZ014573-MZ014617).

### 2.3 Phylogenetic and genetic distance analyses

Previously published sequences of the same genes from cloned reference *T. cruzi* strains from all major DTUs (Breniere et al., 2016; Flores-López and Machado, 2011; Machado and Ayala, 2001, 2002) and from recently sampled US isolates were included in the analyses (S2 Table). *T. cruzi marinkellei* was used as outgroup in the analyses. Sequences were aligned using MUSCLE (Edgar, 2004) and then manually checked. MrBayes 3.1.2 (Huelsenbeck and Ronquist, 2001; Ronquist and Huelsenbeck, 2003) was used to conduct Bayesian analyses using the substitution models chosen by jModeltest on each locus (Posada, 2008). Two independent simultaneous Markov Chain Monte Carlo runs were conducted with four chains each for at least 30,000,000 generations and sampled trees every 10,000 generations. If the standard deviation of split frequencies were not below 0.01 after the first run, the analyses were run for an additional 10,000,000 generations and were stopped after convergence (*i.e.* standard deviation of split frequencies # 0.01). Parameters and corresponding trees were summarized after discarding the initial 25% of each chain as burnin. Maximum likelihood (ML) trees were estimated in PAUP, using the tree bisection-reconnection (TBR) algorithm for the branch swapping (Swofford, 2002b). A reduced data set was selected for the ML analyses due to computational constraints. jModeltest2 was used to estimate the most appropriate nucleotide substitution model used for the ML heuristic search, where the Akaike information criterion (AIC) was employed to select the model (Darriba et al., 2012). Bootstrap support values were obtained by ML analyses of 100 pseudoreplicates of each dataset. Nucleotide sequence differences were calculated in PAUP (Swofford, 2002a) to plot the distribution of pairwise differences within and between DTUs.

### 2.4 Divergence time estimates

A molecular clock was enforced on each data set, and the likelihood of the ML tree was compared to the tree without the clock constraint to determine if any data set did not conform to the molecular clock assumption. The concatenated data set of the 3 nuclear loci (2,757 bp) and the mitochondrial alignments (1,094 bp) were independently used to estimate divergence dates using Bayesian divergence analyses with BEAST 2 (Bouckaert et al., 2014). To calibrate the clock, we used the time of divergence between *T. cruzi* and *T. c. marinkellei* from a previous study (6.23 Mya for the nuclear data set) (Flores-López and Machado, 2011). All other priors and parameters used for this analysis were the same as those previously used (Flores-López and Machado, 2011).

## 3. RESULTS

### 3.1 Phylogenetic diversity of two common *T. cruzi* lineages from Texas and Arizona

We collected *T. cruzi* sequences from fifty-five infected Triatomine bugs from Texas and Arizona (Table S1). Phylogenetic analyses that include published nucleotide sequences from strains representing the 6 currently recognized *T. cruzi* DTUs indicate that the collected isolates are part of two different *T. cruzi* lineages (Figs 2 and 3, S1-S8). 75% of the infected insects carried *T. cruzi* corresponding to TcI (*T. cruzi* DTU I), in agreement with what is known about the genetic diversity of the parasite in North America (Breniere et al., 2016). The other 25% of the infected insects (13/51 from Texas, 3/4 from Arizona) carried isolates that cluster within a well-supported monophyletic group that is more closely related to TcIII-IV-V-VI than to TcI or TcII, but that is significantly divergent from any of the previously defined DTUs.

**Fig. 2.**
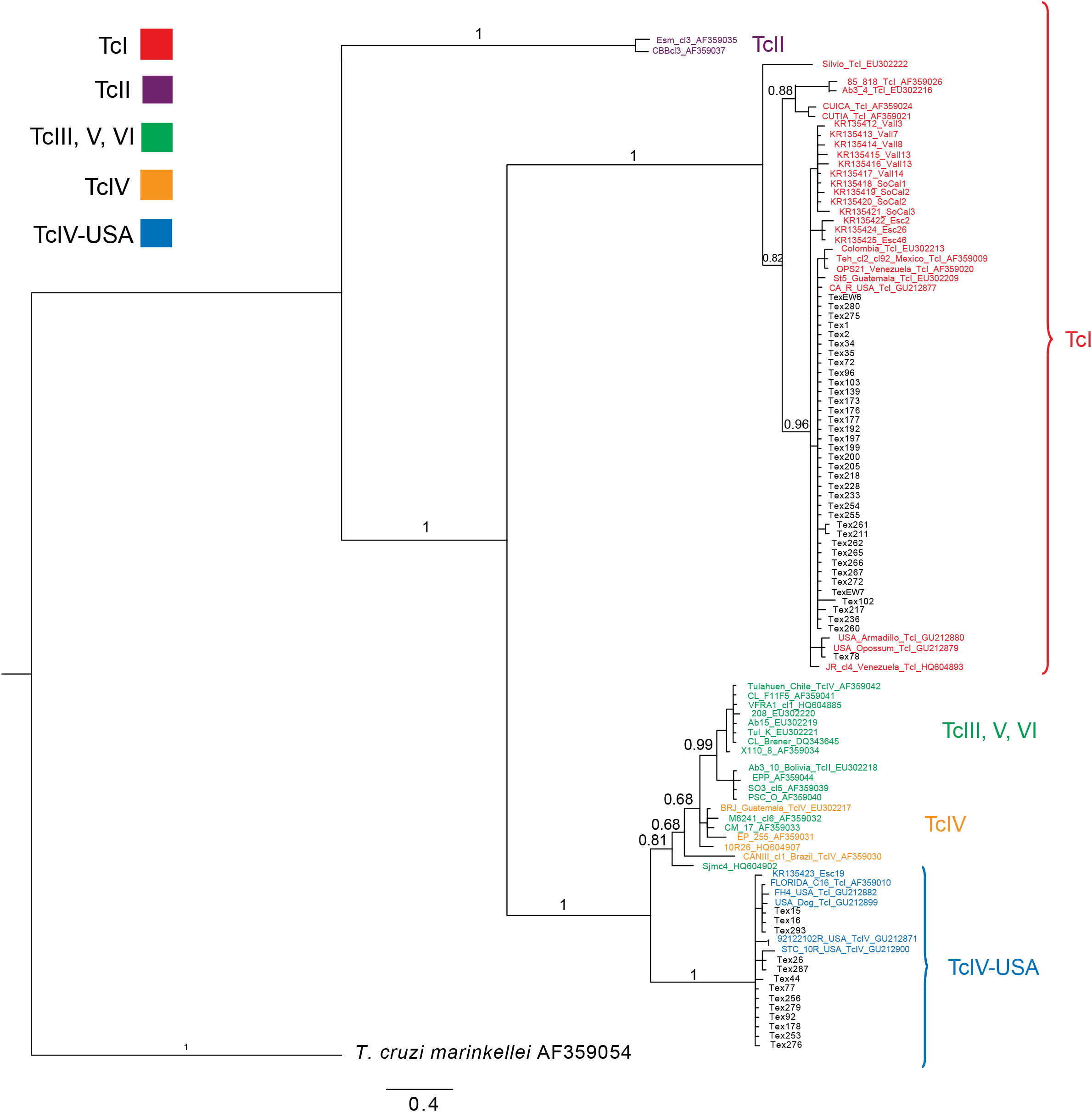
Bayesian phylogenetic tree of the COII-ND1 locus. Reference strains used to represent all previously described *T. cruzi* DTU’s have Genbank accession numbers attached to the strain ID. Original sources of reference strains are listed in Table S2. Texas samples from this study are noted in black font. Values above internal nodes represent posterior probability support values. Scale bar represents the number of substitutions per site. *T. cruzi marinkellei* was used an outgroup in the rooting of the tree.

**Fig. 3.**
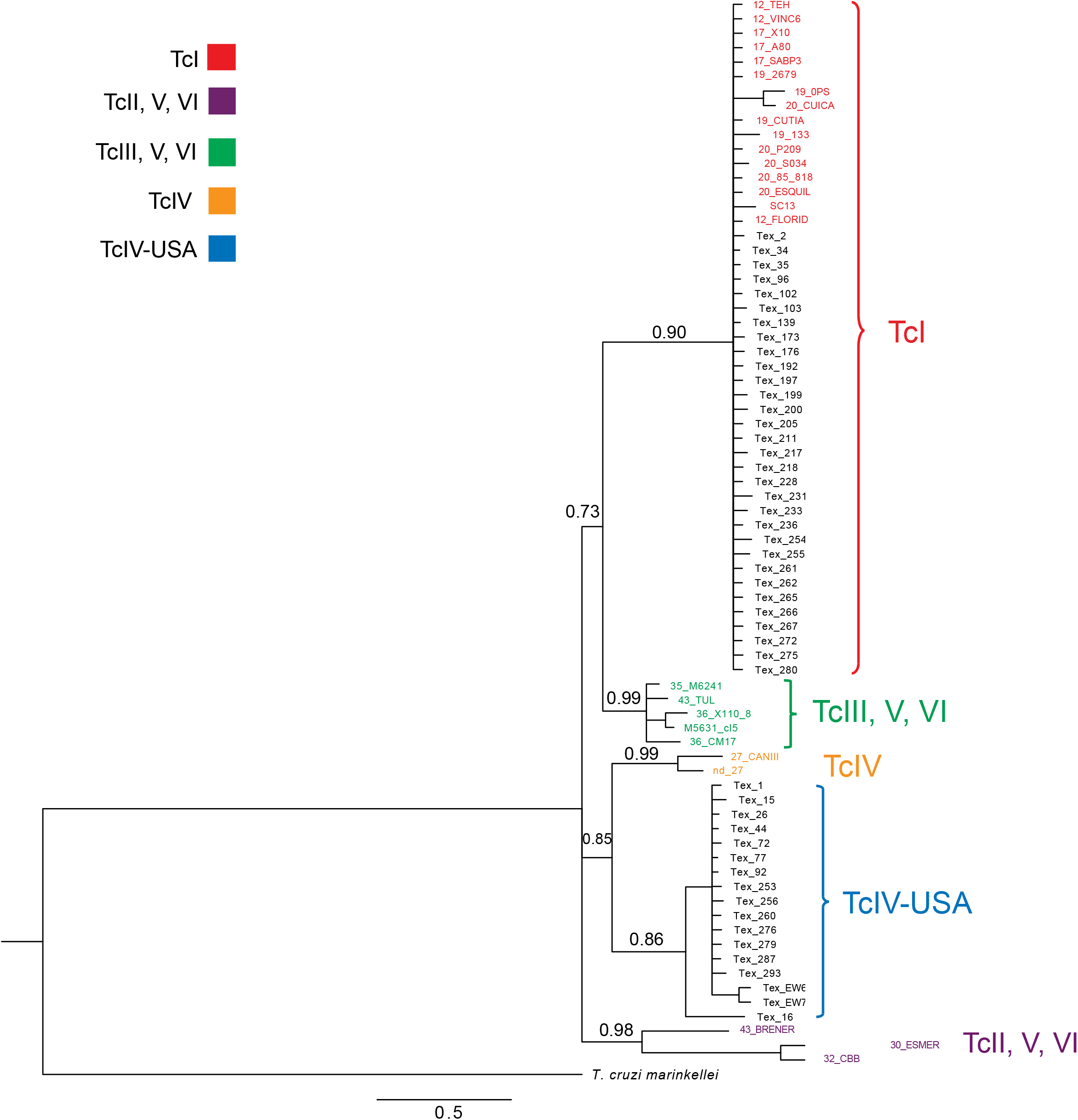
Bayesian phylogeny of concatenated nuclear data set. The total data set consists of 2,757 nucleotides from 3 markers (TcCLB.506529.310, MSH2, DHFR-TS). Texas samples from this study are noted in black font. Original sources of reference strains are listed in Table S2. Values above internal nodes represent posterior probability support values. Scale bar represents the number of substitutions per site. *T. cruzi marinkellei* was used an outgroup in the rooting of the tree. Sequences from DTU’s V and VI cluster in two separate clades of the phylogeny due to their hybrid nature (Machado and Ayala, 2001).

Given that the vast majority of DTU reference sequences are from South American isolates, we surveyed studies from the USA to incorporate all published sequences from the same loci used in our analyses. We found that the non-TcI lineage from our survey corresponds to what had been previously described as TcIV from North America (Barnabe et al., 2001; Lewis et al., 2009; Marcili et al., 2009). This lineage is closer to TcIV when considering nuclear loci but is equally divergent to TcIII-IV-V-VI at the mitochondrial locus (Figs 2, 3, S1-S8). Based on our survey of GenBank and the literature, no sequences that cluster with South American TcIV have been collected in the USA. The closest sequences to this non-TcI lineage from our survey come from isolates that have been termed either TcIV (acknowledging that there is sequence divergence between South and North America isolates) (Messenger et al., 2012; Roellig et al., 2013; Yeo et al., 2011), TcIV-US (Lewis et al., 2011) or TcIV-USA (Shender et al., 2016) which is the label we will use in this report. All previously published sequences (available in GenBank) that cluster with the TcIV-USA isolates from our survey are included in all the phylogenies presented here (Figs 2, 3, S1-S8; Table S2) as well as in the distance analyses presented below in section 3.2. Interestingly, TcIV-USA isolates were detected in all the five North American triatomine species sampled in this study (*T. gerstaeckeri, T. lecticularia, T.indictiva, T. sanguisuga* and *T. recurva*).

Sequences from all TcIV-USA isolates are reciprocally monophyletic regardless of the locus, the combination of loci, or method used (Figs 2, 3, S1-S8). While DTUs TcIII, TcV and TcVI are effectively indistinguishable from each other with the sequences used, TcIV-USA is clearly different from all other DTUs (Figs 2, 3, S1-S8), including DTU TcIV which is represented by the only two reference sequences available from two South American isolates (CANIIIcl1, EPP). Bayesian posterior probability values indicate high support for this monophyletic group regardless of the dataset used for the analyses (Figs 2, 3, S1-S5). ML bootstrap support values are also high for this clade regardless of the dataset (S6-8 Figs). In the analyses of the mtDNA data (COII-NDI), TcIV-USA sequences form a monophyletic group whose closest relative is a monophyletic group composed by TcIII-IV-V-VI (Figs 2, S1, S6). This same set of relationships are recovered in analyses of the concatenated data from all the 4 loci (Figs S5, S8). In the analyses of concatenated sequences from the 3 nuclear loci (DHFR-TS, MSH2, TcCLB.506529.310), TcIV-USA and TcIV sequences cluster together albeit with evidence of significant divergence between the two clades (Figs 3, S7, and see Section 3.2). Further, focusing on the nuclear loci, the TcIV-TcIV-USA clade appears to be well supported only in locus DHFR-TS (Fig. S4).

Estimates of the divergence time of TcIV-USA from its common recent ancestor with South American TcIV is similar for both nuclear (110,000 yrs ago) and mitochondrial data (120,000-160,000 yrs ago) (Table S3), indicating a mid to late Pleistocene divergence and entry into North America, well before the arrival of humans into North America.

### 3.2 Nucleotide divergence between TcIV-USA and other *T. cruzi* DTUs

To complement our phylogenetic analyses, we plotted the distribution of nucleotide pairwise differences within and between DTUs to determine if TcIV-USA is significantly divergent from other DTUs and could thus correspond to an independent evolutionary lineage. This approach looks for a “barcoding gap” to delimit evolutionary lineages comparing the distribution of pairwise nucleotide differences within and between lineages (Candek and Kuntner, 2015; Hebert et al., 2004; Meyer and Paulay, 2005), and here we use the previously delimited DTUs as controls. We found that the distribution of intra-DTU sequence divergence is significantly lower than the distribution of inter-DTU sequence divergence for any of the comparisons between the TcIV-USA isolates and any of the six previously defined DTUs that are either distantly (TcI, TcII) or more closely related (TcIII, TcIV, TcV, TcVI) (Figs 4, S9, S10). All the comparisons of intra vs inter-DTU genetic distances that include the TcIV-USA isolates are statistically significant (Mann Whitney U-tests: p<0.00001). Furthermore, the divergence between the TcIV-USA isolates and its four most closely related DTUs (TcIII, TcIV, TcV, TcVI) is significantly higher than the level of divergence among those four DTUs (Figs 4, S9, S10). In fact, DTUs TcIII, TcV and TcVI are effectively indistinguishable from each other using the four surveyed genes. Moreover, if one were to force TcIV and TcIV-USA sequences into a single genetic group, the intraclonal nucleotide sequence diversity of this forced TcIV/TcIV-USA group would be significantly higher than the diversity of any of the previously defined DTUs (Fig. 5). All these results indicate that TcIV-USA is significantly divergent from other *T. cruzi* DTUs and that this finding is not caused by the paucity of TcIV sequences from South American samples.

**Fig. 4.**
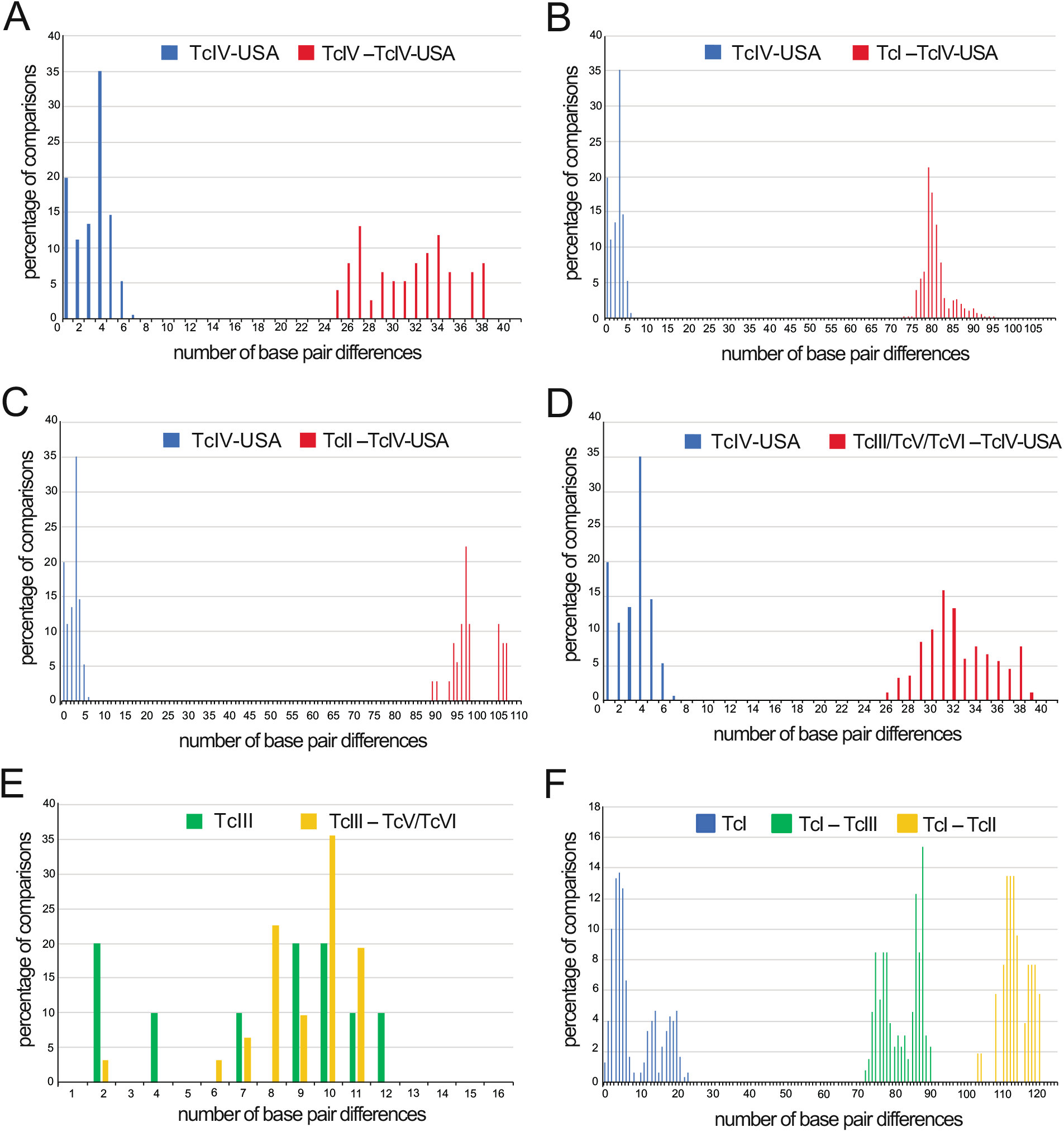
Distribution of intra and inter-DTU genetic distances for COII-NDI. Absolute pairwise nucleotide differences were calculated within and between DTU’s. Sources of the sequences used for each DTU are listed in Table S2. Differences between intra and inter-DTU differences were analyzed using Mann-whitney U-tests. **A**: TcIV—TcIV-USA (p<0.0001); **B**: TcI—TcIV-USA (p<0.0001); **C**: TcII—TcIV-USA (p<0.0001); **D**: TcIII/V/VITcIV—USA (p<0.0001); **E**: TcIII— TcV/VI (p=0.42372, N.S); **F**: TcI—TcIII (p<0.00001), TcI—TcII (p<0.00001). p-values were adjusted for multiple tests using the Bonferronni correction.

**Fig. 5.**
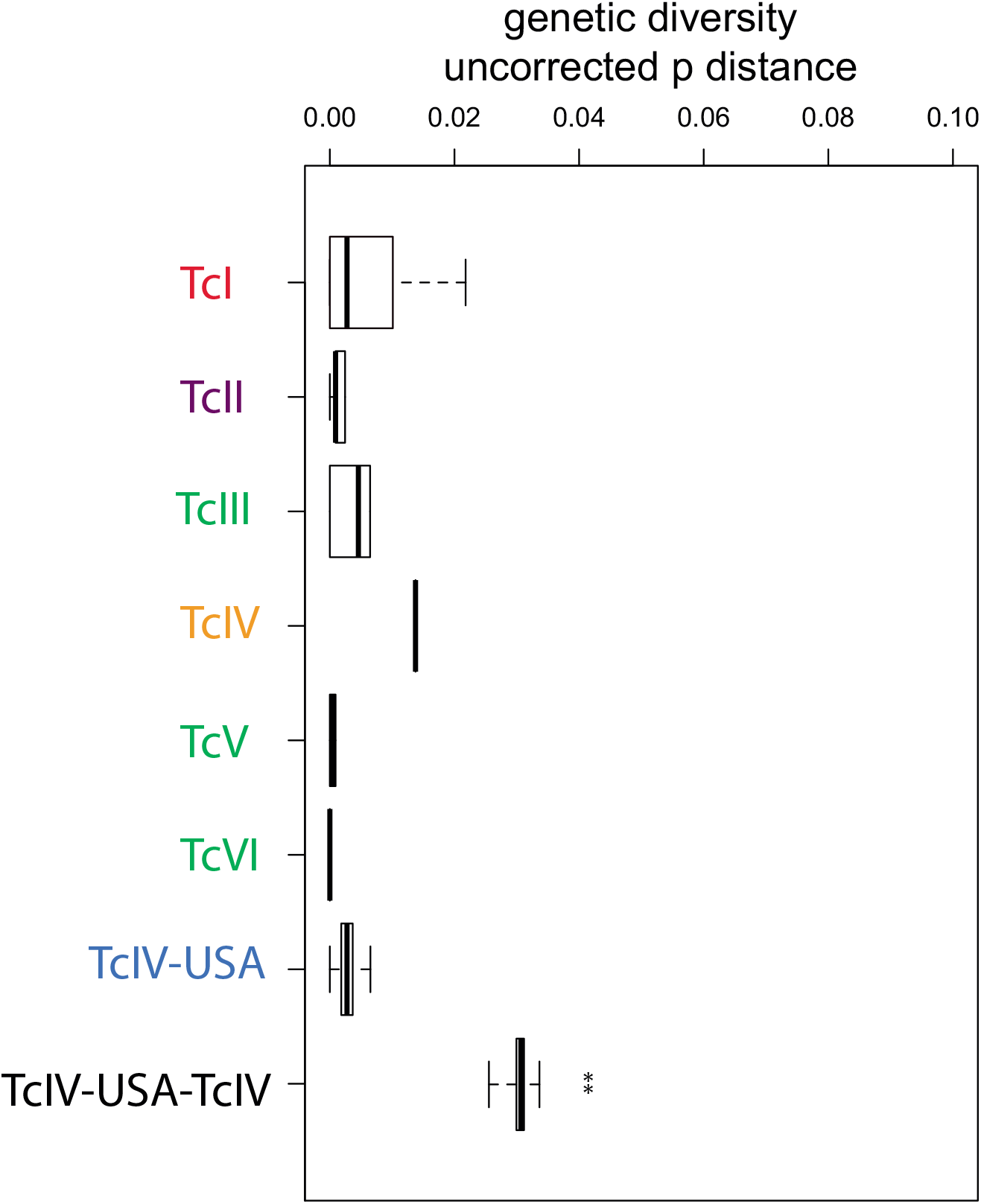
Levels of intra-DTU nucleotide diversity in COII-ND1. Labels in the X-axis indicate the various *T cruzi* DTUs. The level of nucleotide diversity observed when combining TcIV and TcIV-USA is significantly higher than that of any of the other DTUs (A-D: p < 2e-16; E: p=1; F: p < 2e-16; Wilcoxon signed-rank test, with Bonferroni adjusted p-values).

### 3.3 Evidence of genetic exchange events in TcIV-USA

Genetic exchange between different *T. cruzi* lineages can generate phylogenetic incongruency among different genetic markers (Machado and Ayala, 2001, 2002). A *T. cruzi* isolate from a *T. sanguisuga* specimen collected in Gainesville (Florida_C16) (Barnabe et al., 2000) provided the first evidence of mitochondrial introgression in this parasite based on phylogenetic incongruency between markers (Machado and Ayala, 2001). This sample’s COII-ND1 sequence actually represents the first published sequence of TcIV-USA (Figs 2, S2, S6) (although that label was not used at the time), while its sequences from nuclear genes DHFR and TR cluster with TcI (Machado and Ayala, 2001, 2002) in agreement with its DTU classification (TcI) based on allozyme data (Barnabe et al., 2000). These findings strongly suggested that Florida_C16 represented a strain that arose from genetic exchange between TcI and an unknown lineage (at that time) that we now know is TcIV-USA. In that instance of inferred genetic exchange, the TcIV-USA mitochondria (maxicircle) introgressed into a TcI nuclear genetic background.

In our study there are three samples from Texas (Tex1, Tex72, Tex260) that also have conflicting phylogenies between loci. COII-ND1 sequences place all of them in TcI (Figs 2, S1), while sequences from nuclear loci TcCLB.506529.310 (Fig. S2) and MSH2 (Fig. S3) place them in TcIV-USA (DHFR-TS was not sequenced in these samples). Most nuclear loci in the two major hybrid strains of *T. cruzi* (TcV & TVI) that have been cloned and maintained in culture have usually multiple heterozygous sites due to mixed parental ancestry (TcII and TcIII) of their nuclear DNA (Flores-López and Machado, 2011; Machado and Ayala, 2001, 2002). Since the mitochondria are thought to have a uniparental mode of inheritance, the presence of heterozygous sites in a mitochondrial chromatogram indicates mixed amplification of two distinct genotypes and is direct evidence of a mixed infection. None of those three Texas samples (Tex1, Tex72, Tex260) show ambiguous sites in their mitochondrial chromatograms, but sample Tex72 shows ambiguous sites in the MSH2 loci at all the nucleotide sites that distinguish TcI from TcIV-USA (Fig. S11) suggesting this isolate is heterozygous for TcI and TcIV-USA alleles consistent with a potential recent hybridization event.

## 4. DISCUSSION

We collected nucleotide sequences from three single-copy nuclear genes (DHFR-TS, MSH2, TcCLB.506529.310) and one mitochondrial region (COII-NDI) to characterize *T. cruzi* genetic variation in infected Triatomines from several Texas and Arizona locations. Nucleotide sequences from multiple loci provide more resolution to identify DTUs and uncover phylogenetic relationships than data from single, short or less variable regions like those traditionally used in large field surveys. We found two genetic lineages of *T. cruzi* circulating in the sampled Triatomines. DTU TcI was the most common lineage, found in 75% of the infected triatomines (n=55). The second lineage, found in 25% of the infected triatomines, showed significant genetic divergence from previously defined *T. cruzi* DTUs, but corresponds to a previously described divergent North American branch of DTU TcIV labelled TcIV-USA or TcIV-US (Lewis et al., 2011; Messenger et al., 2012; Roellig et al., 2013; Shender et al., 2016; Yeo et al., 2011). Using evidence from phylogenetic and genetic distance analyses, our results indicate that TcIV-USA is significantly divergent from TcIV and from any other *T. cruzi* DTUs.

The divergence between South American and North American TcIV has been noticed in previous studies based on 1) multilocus enzyme electrophoresis (MLEE) (Barnabe et al., 2001); 2) RFLP and PCR fragment size differences (Lewis et al., 2009), and; 3) nucleotide sequences from SSU rRNA and Cyt b (Marcili et al., 2009). More recently, this divergence was also reported based on phylogenetic analyses of North American samples using sequences from few loci (Lewis et al., 2011; Messenger et al., 2012; Roellig et al., 2013; Shender et al., 2016; Yeo et al., 2011). Interestingly, this South America-North America TcIV divergence was even noticeable in the MLEE and RAPD cladograms used to define the currently accepted six *T. cruzi* DTUs (Brisse et al., 2000), although in that study the North American TcIV was represented by a single North American isolate (DogTheis). Based on the observed level of RAPD and MLEE divergence between South American TcIV isolates and the only North America TcIV isolate, plus the strong congruence between cladograms (see clade “2a” from Figure 3 of (Brisse et al., 2000)), it is worth wondering whether that North American TcIV lineage would have been described as a separate DTU had the Brisse et al study (Brisse et al., 2000) included more isolates from North America. Our phylogenetic analyses confirm the divergence of TcIV-USA, and our genetic distance analyses (Figs 4, 5, S9, S10) demonstrate that this divergence is significant relative to all other defined DTUs. Based on our survey of the literature and GenBank, no sequences similar to the South American TcIV DTU have been collected in the USA. Future efforts using genomic data should allow a rigorous investigation of genome-wide levels of divergence among all DTUs, and should determine if any changes in the placement of TcIV-USA within the current *T. cruzi* classification system are warranted.

The geographic distribution and host associations of TcIV-USA (“TcIV” as mostly used in US-based studies) are quite broad, covering for the most part every state, mammalian host and triatomine species where *T. cruzi* has been detected. In our study, TcIV-USA was found in all the five triatomine species sampled: four species from Texas (*T. gerstaeckeri, T. lecticularia, T.indictiva, T. sanguisuga)* and one species from Arizona (*T. recurva*). Our sample sizes are not large enough to conduct statistical tests of preferred vector-DTU associations, but recent work (Curtis-Robles et al., 2017) reported that in Texas *T. gerstaeckeri* was disproportionately associated with TcI while *T. sanguisuga* was mostly associated with TcIV-USA (identified as “TcIV” in that study). Although our data from Arizona were obtained from a small number of *T. recurva* infected specimens, it is important to note that, by far, *T. rubida* is the most abundant species in Arizona. *T. rubida* has repeatedly been found infected with *T. cruzi* (Reisenman et al., 2010; Reisenman et al., 2012) although DTUs were not identified. In the neighboring state of New Mexico, a recent study found a small proportion of *T. rubida* infected with TcIV, and even one specimen with a mixed TcI-TcIV infection (Rodriguez et al., 2021). Furthermore, the second most abundant Triatomine species in Arizona is *T. protracta,* also not sampled in this study, which happens to be the most common species in California. A recent study reported the presence of TcIV-USA in California associated with *T. protracta* (Shender et al., 2016). Interestingly, that study uncovered *T. protracta* infected with “TcIV” but only in Southern California, while TcI was present in all areas of the state that were sampled, although sample sizes were small.

The most comprehensive USA survey of Triatomines infected with *T. cruzi* with a focus in Texas and few samples from 16 southern US states (Curtis-Robles et al., 2018), reported high frequencies of infection with TcI and “TcIV” in Texas in the same four *Triatoma* species we sampled (*T. gerstaeckeri, T. lecticularia, T.indictiva, T. sanguisuga*). In addition, “TcIV” was observed in eight states but only associated with *T. sanguisuga*, although samples sizes were very small. Although in that survey (Curtis-Robles et al., 2018) phylogenies were not presented to support DTU identification, we assume that samples identified as “TcIV” are TcIV-USA indicating that this lineage is commonly associated with Triatomines and widespread in southern USA. Additional surveys should be conducted to determine the most common mammals associated with TcIV-USA. Additionally, a sample isolated from a *T. dimidiata* specimen collected from the Yucatan peninsula in southern Mexico also clusters within TcIV-USA (CAFL unpublished). Future work should incorporate variable sequence markers like those used here to confirm the identity and frequency of the *T. cruzi* lineages circulating in Triatomine insects in the USA.

The observation of multiple instances of potential TcI-TcIV-USA hybrids (TcI-TcIV in the literature) (Lewis et al., 2011; Machado and Ayala, 2001; Messenger et al., 2012; Roellig et al., 2013; Shender et al., 2016) is not surprising given that mixed infections in vectors and mammal hosts are routinely reported (Curtis-Robles et al., 2018; Curtis-Robles et al., 2016; Herrera et al., 2015; Rodriguez et al., 2021; Roellig et al., 2008), given that these are the most common lineages in the USA (Breniere et al., 2016), and given that TcI and TcIV-USA have been coexisting in North America probably for more than 100,000 years (Table S3). A *T. cruzi* sample that was isolated from a *T. sanguisuga* specimen collected in Gainesville (Florida_C16) and that has been classified as TcI based on allozyme data (Barnabe et al., 2000), clustered with strong support within the TcIV-USA clade in the mitochondrial analysis (Figs 2, S1) but with TcI in the analyses of the concatenated nuclear loci (Fig. 3). That incongruent phylogenetic placement among loci is consistent with hybridization/introgression between TcI and TcIV-USA. This hybridization event was previously described two decades ago, but since there were no published sequences from TcIV-USA at that time, the mtDNA sequence was basally positioned in the TcIII clade (Machado and Ayala, 2001). A similar pattern of phylogenetic incongruency (mtDNA: TcIV-USA, nuclear loci: TcI) has been reported for 1 isolate from Georgia (Messenger et al., 2012), and for nine isolates from various USA states (Roellig et al., 2013). Here we report the opposite pattern of phylogenetic incongruency (mtDNA: TcI, nuclear loci: TcIV-USA) in three samples from Texas (Tex1, Tex72 & Tex260), consistent with previous reports from few isolates of *T. protracta* in California (Shender et al., 2016) and from a dog isolate from Tennessee (Roellig et al., 2013).

Our findings also confirm previous studies indicating that TcI is the most common lineage of *T. cruzi* in the USA (Barnabe et al., 2001; Breniere et al., 2016; Curtis-Robles et al., 2018; Curtis-Robles et al., 2017; Garcia et al., 2017; Herrera et al., 2015; Meyers et al., 2017; Roellig et al., 2008; Roellig et al., 2013; Shender et al., 2016; Vandermark et al., 2018). This is not surprising, given that TcI is also the most frequently reported lineage found in neighboring Mexico and Central America (Dorn et al., 2017; Zumaya-Estrada et al., 2012). Phylogenetic studies that incorporate the large genetic diversity of TcI found across the Americas showed that TcI samples isolated from North America tend to form a distinct cluster nested within the TcI isolates from South America (Cura et al., 2010; Llewellyn et al., 2009). This suggests that the TcI lineages found in North America may be derived from South America, although the timing of the North American introduction is unclear. Although some authors suggested that the dispersal could have been facilitated by humans (Lewis et al., 2011), our divergence time estimates indicate that the origin of the North American TcI clade greatly predates humans’ presence in the Americas (Table S3).

Although Chagas disease is relatively uncommon in the USA, a small but increasing number of autochthonous vectorial transmission cases have been reported in the last decade in southern states. Environmental changes and globalization will undoubtedly increase the opportunities for vectorial transmission of *T. cruzi* in the USA (Eberhard et al., 2020; Garza et al., 2014), and future efforts should thus focus on evaluating the epidemiological importance of TcI and TcIV-USA. Our study emphasizes the need for appropriate epidemiological surveillance and vector control programs in southern USA states. Infection rates, lack of domiciliated vectors, socioeconomic factors and habitat conditions that are usually associated with Chagas disease in Mexico, Central and South America are much less common in the USA. Nevertheless, the epidemiological importance and prevalence of TcIV-USA merits investigation, as well as its geographical range and pathogenicity in humans.

### 4.1 Conclusions

This study confirms that two major genetic lineages of *T. cruzi* (TcI, TcIV-USA) circulate in the Southwest USA. Parasite sequences belonging to TcIV-USA have been previously reported in other studies from the USA as “TcIV”. However, our phylogenetic analyses and a barcoding gap approach show that sequences from this lineage are significantly different to those from South American TcIV and from those of any other currently defined DTUs. A whole-genome comparison approach will be needed to properly determine the most appropriate taxonomic positioning of this North American genetic lineage within *T. cruzi*. Although the correlation between the variable clinical symptoms of Chagas disease and the genetic background of *T. cruzi* is not yet fully understood (Jimenez et al., 2019), having a better understanding of the genetic variability of *T. cruzi* in the USA and the potential connection between genetic variants and disease progression will be of major importance. Thus, knowledge of the identity of *T. cruzi* lineages circulating in the USA and their possible connections with human infections and disease is important for designing potential treatments. In fact, a more vigilant approach to study clinical manifestations of Chagas in the context of *T. cruzi* DTU identity is now urgently needed in the USA.

## AKNOWLEDGEMENTS

We thank John Hildebrand for his support in many aspects of this work, and to Teresa Gregory, William Savary and Blake Sissel for help collecting and identifying insects. We would also like to thank Peggy Billingsley for her contributions to the molecular portion of this work. Sequencing supported by funds from the University of Maryland to C.A.M.

## Supporting Information

**Table S1.** Geographical collection information for the samples from Texas and Arizona.

**Table S2.** Additional published sequences used in the analyses. * Accession numbers that were not included in the figures, but confirmed to be TcIV-USA in phylogenetic analyses (not shown).

**Table S3**. Bayesian estimates of divergence (time in millions of years) for the main *T. cruzi* lineages and TcIV-USA.

**Fig. S1. Bayesian phylogenetic tree of COII-ND1 data set with TcDOM reference strains.**

Data set consists of a 417 bp fragment from the cytochrome oxidase subunit II and NADH dehydrogenase subunit 1 loci (COII-ND1). Genbank Accesion numbers JX431235-JX431259 were reported in (Zumaya-Estrada et al., 2012) as TcDOM. Values above internal nodes represent posterior probability support values. Scale bar represents the number of substitutions per site.

**Fig. S2. Bayesian phylogenetic tree of the TcCLB.506529.310 nuclear locus.** All major *T. cruzi* clades (TcI-VI) and new *T. cruzi* lineage (TcIV-USA) are highlighted in the phylogeny (except TcBAT). This was the only locus sequenced from the Arizona samples. Values above internal nodes represent posterior probability support values. Scale bar represents the number of substitutions per site. Sequences from DTU’s V and VI cluster in two separate clades of the phylogeny due to their hybrid nature (Machado and Ayala, 2001).

**Fig. S3. Bayesian phylogenetic tree of the MSH2 sequence data.** All major *T. cruzi* clades (TcI-VI) are highlighted in the phylogeny. NCBI accesion codes GU212992-GU213034 are from Roellig et al (Roellig et al., 2013). Scale bar represents number of substitutions per site. Sequences from DTU’s V and VI cluster in two separate clades of the phylogeny due to their hybrid nature (Machado and Ayala, 2001).

**Fig. S4. Bayesian phylogenetic tree of the DHFR sequence data.** All major *T. cruzi* clades (TcI-VI) are highlighted in phylogeny (except TcBat). Values above internal nodes represent posterior probability support values. Scale bar represents number of substitutions per site.

**Fig. S5. Bayesian phylogeny of concatenated data set.** The total data set consists of 3,851 nucleotides from all 4 markers used in this study (COII-ND1, TcCLB.506529.310, MSH2, DHFR-TS). Texas samples from this study are noted in black font. Original sources of reference strains are listed in S2 Table. Values above internal nodes represent posterior probability support values. Scale bar represents number of substitutions per site.

**Fig. S6. Maximum likelihood phylogenetic tree of the COII-ND1 locus.** The data set included a total of 1,229 nucleotides. Values above nodes indicate bootstrap support values. The nucleotide substitution model used was the TrN+G, selected by the AIC.

**Fig. S7. Maximum likelihood phylogenetic tree of the concatenated data set of 3 nuclear markers.** The data set included a total of 2,757 nucleotides. Values above nodes indicate bootstrap support values. The nucleotide substitution model used was the TPM1uf+G, selected by the AIC.

**Fig. S8. Maximum likelihood phylogenetic tree of the concatenated data set of both mitochondrial and nuclear markers.** The data set included a total of 3,851 nucleotides including all 4 loci sampled in this study. Values above nodes indicate bootstrap support values. The nucleotide substitution model used was the TVM+G, selected by the AIC.

**Fig. S9. Distribution of intra and inter-DTU genetic distances for MSH2.** Absolute pairwise nucleotide differences were calculated within and between DTU’s. Sequences used for each DTU can be seen in S2 Table. The significance of the differences between intra and inter-DTU differences were obtained using the Mann-whitney U-test. **A**: TcIV—TcIV-USA (p<0.00001); **B**: TcI—TcIV-USA (p<0.00001); **C**: TcII—TcIV-USA (p<0.00001); **D**: TcII/TcV/TcVI—TcIV-USA (p<0.00001); **E**: TcI—TcII (p<0.00001); **F**: TcII—TcIII/TcV/TcVI (p<0.00001).

**Fig. S10. Distribution of intra and inter-DTU genetic distances for DHFR-TS.** Absolute pairwise nucleotide differences were calculated within and between DTU’s. Sequences used for each DTU can be seen in S2 Table. The significance of the differences between intra and inter-DTU differences were obtained using the Mann-whitney U-test. **A**: TcIV—TcIV-USA (p<0.00001); **B**: TcI—TcIV-USA (p<0.00001); **C**: TcII—TcIV-USA (p<0.00001); **D**: TcIII/TcV/TcVI—TcIV-USA (p<0.00001); **E**: TcIII—TcV/TcVI (p=0.1615); F: TcI—TcIII (p<0.00001), TcI—TcII (p<0.00001).

**Fig. S11. Chromatograms showing location of ambiguous sites in the MSH sequence of sample Tex72.**

## Notes

### Competing Interest Statement

The authors have declared no competing interest.

