## Supplemental Tables and Figures for "Phylogenetic diversity of two common *Trypanosoma cruzi* lineages in the Southwestern United States"

**Suplementary Table 1.** Geographical location of the isolated samples from Texas and Arizona.

| **Sample ID** | **Genus** | **Species** | **Longitude** | **Latitude** | **Location** |
|---|---|---|---|---|---|
| Tex1 | *Triatoma* | *gerstaeckeri* | -99.250 | 29.56812 | Hondo, Texas |
| Tex2 | *Triatoma* | *gerstaeckeri* | -99.250 | 29.56812 | Hondo, Texas |
| Tex15 | *Triatoma* | *sanguisuga* | -97.375 | 30.217 | Elgin, Texas |
| Tex16 | *Triatoma* | *sanguisuga* | -97.375 | 30.217 | Elgin, Texas |
| Tex26 | *Triatoma* | *indictiva* | -98.186 | 30.235 | Hays County, Texas |
| Tex34 | *Triatoma* | *gerstaeckeri* | -99.766 | 29.349 | Uvalde County, Texas |
| Tex35 | *Triatoma* | *gerstaeckeri* | -99.766 | 29.349 | Uvalde County, Texas |
| Tex44 | *Triatoma* | *gerstaeckeri* | -99.766 | 29.349 | Uvalde County, Texas |
| Tex72 | *Triatoma* | *lecticularia* | -99.179314 | 29.18498541 | Medina County, Texas |
| Tex77 | *Triatoma* | *gerstaeckeri* | -99.179314 | 29.18498541 | Medina County, Texas |
| Tex78 | *Triatoma* | *gerstaeckeri* | -99.179314 | 29.18498541 | Medina County, Texas |
| Tex92 | *Triatoma* | *gerstaeckeri* | -99.179314 | 29.18498541 | Medina County, Texas |
| Tex96 | *Triatoma* | *gerstaeckeri* | -99.179314 | 29.18498541 | Medina County, Texas |
| Tex102 | *Triatoma* | *gerstaeckeri* | -99.179314 | 29.18498541 | Medina County, Texas |
| Tex103 | *Triatoma* | *gerstaeckeri* | -99.179314 | 29.18498541 | Medina County, Texas |
| Tex139 | *Triatoma* | *gerstaeckeri* | 98.2548052 | 26.09747629 | 519 8th St., Hidalgo, Texas, 78557 |
| Tex173 | *Triatoma* | *gerstaeckeri* | -97.227244 | 26.073822 | 02 E.A.C.P. 1765 hwy 100, Port Isabel, TX |
| Tex176 | *Triatoma* | *gerstaeckeri* | -98.070297 | 26.530892 | 03 E.Q. La Sal del Rey, Hidalgo County, Texas |
| Tex177 | *Triatoma* | *gerstaeckeri* | -98.070297 | 26.530892 | 4 E.Q. La Sal del Rey, Hidalgo County, Texas |
| Tex178 | *Triatoma* | *gerstaeckeri* | -98.344465 | 26.398811 | 03 E.A.C.P. 22675 N. Moorefield Rd., Edinburg, TX, 78541 |
| Tex192 | *Triatoma* | *gerstaeckeri* | -98.350567 | 26.181283 | Hidalgo County TX |
| Tex197 | *Triatoma* | *gerstaeckeri* | -98.350567 | 26.181283 | Hidalgo County TX |
| Tex199 | *Triatoma* | *gerstaeckeri* | -98.350567 | 26.181283 | Hidalgo County TX |
| Tex200 | *Triatoma* | *gerstaeckeri* | -98.350567 | 26.181283 | Hidalgo County TX |
| Tex205 | *Triatoma* | *gerstaeckeri* | -98.350567 | 26.181283 | Hidalgo County TX |
| Tex211 | *Triatoma* | *gerstaeckeri* | -98.350567 | 26.181283 | Hidalgo County TX |
| Tex217 | *Triatoma* | *gerstaeckeri* | -98.350567 | 26.181283 | Hidalgo County TX |

| | | | | | |
|---|---|---|---|---|---|
| Tex218 | *Triatoma* | *gerstaeckeri* | -98.350567 | 26.181283 | Hidalgo County TX |
| Tex228 | *Triatoma* | *gerstaeckeri* | -98.350567 | 26.181283 | Hidalgo County TX |
| Tex231 | *Triatoma* | *gerstaeckeri* | -98.350567 | 26.1813 | Hidalgo County TX |
| Tex233 | *Triatoma* | *gerstaeckeri* | -98.350567 | 26.1813 | Hidalgo County TX |
| Tex236 | *Triatoma* | *gerstaeckeri* | -98.350567 | 26.1813 | Hidalgo County TX |
| Tex253 | *Triatoma* | *lecticularia* | -99.17651 | 28.58779 | La Salle Cty, TX |
| Tex254 | *Triatoma* | *lecticularia* | -99.17651 | 28.58779 | La Salle Cty, TX |
| Tex255 | *Triatoma* | *gerstaeckeri* | -99.177253 | 28.58596 | La Salle Cty, TX |
| Tex256 | *Triatoma* | *gerstaeckeri* | -99.177253 | 28.58596 | La Salle Cty, TX |
| Tex260 | *Triatoma* | *lecticularia* | -99.250555 | 29.578333 | Medina County, Texas |
| Tex261 | *Triatoma* | *gerstaeckeri* | -99.250555 | 29.578333 | Medina County, Texas |
| Tex262 | *Triatoma* | *gerstaeckeri* | 99.2505556 | 29.586667 | Medina County, Eagle Bluff Ranch, TX |
| Tex265 | *Triatoma* | *gerstaeckeri* | -98.35055 | 26.18121667 | Bastrop Cty, TX |
| Tex266 | *Triatoma* | *gerstaeckeri* | -98.35055 | 26.18121667 | Bastrop Cty, TX |
| Tex267 | *Triatoma* | *gerstaeckeri* | -98.35055 | 26.18121667 | Bastrop Cty, TX |
| Tex272 | *Triatoma* | *gerstaeckeri* | -98.35055 | 26.18121667 | Bastrop Cty, TX |
| Tex275 | *Triatoma* | *gerstaeckeri* | -98.35055 | 26.18121667 | Bastrop Cty, TX |
| Tex276 | *Triatoma* | *gerstaeckeri* | -98.35055 | 26.1812167 | Bastrop Cty, TX |
| Tex279 | *Triatoma* | *gerstaeckeri* | -98.35055 | 26.18121667 | Bastrop Cty, TX |
| Tex280 | *Triatoma* | *gerstaeckeri* | -98.35055 | 26.18121667 | Bastrop Cty, TX |
| Tex287 | *Triatoma* | *gerstaeckeri* | -97.9563 | 30.146613 | 24 Concord Circle, Austin, Hays County, TX, 78737 |
| Tex293 | *Triatoma* | *gerstaeckeri* | -97.570179 | 30.102131 | Bastrop Cty, TX |
| EW6 | *Triatoma* | *gerstaeckeri* | No data | No data | Uvalde County, Texas |
| EW7 | *Triatoma* | *gerstaeckeri* | No data | No data | Uvalde County, Texas |
| DF78 (09-371) | *Triatoma* | *recurva* | No data | No data | Rio Rico, AZ |
| DF79 (09-252) | *Triatoma* | *recurva* | No data | No data | Vail, AZ |
| DF80 (09-502) | *Triatoma* | *recurva* | No data | No data | Rio Rico, AZ |
| DF81 (09-160) | *Triatoma* | *recurva* | No data | No data | Tubac, AZ |

**Supplementary Table 2.** Additional sequences used in the analyses. Accession numbers with * were not included in the figures but confirmed to be TcIV-USA in phylogenetic analyses (not shown).

| Strain/isolate | GenBank Accession # | Molecular marker | DTU (from study) | Source |
|---|---|---|---|---|
| Esmeraldo cl3 | AF359035 | COII-NDI | TcII | [1] |
| CBB cl3 | AF359037 | COII-NDI | TcII | [1] |
| Silvio X10 cl.4 | EU302222 | COII-NDI | TcI | [2] |
| 85/818 | AF359026 | COII-NDI | TcI | [1] |
| Ab3-4 | EU302216 | COII-NDI | TcI | [2] |
| CUICA cl1 | AF359024 | COII-NDI | TcI | [1] |
| CUTIA cl1 | AF359021 | COII-NDI | TcI | [1] |
| Vall3 | KR135412 | COII-NDI | TcI | [3] |
| Vall7 | KR135413 | COII-NDI | TcI | [3] |
| Vall8 | KR135414 | COII-NDI | TcI | [3] |
| Vall13 | KR135415 | COII-NDI | TcI | [3] |
| Vall13 | KR135416 | COII-NDI | TcI | [3] |
| Vall14 | KR135417 | COII-NDI | TcI | [3] |
| SoCal1 | KR135418 | COII-NDI | TcI | [3] |
| SoCal2 | KR135419 | COII-NDI | TcI | [3] |
| SoCal2 clone 2 | KR135420 | COII-NDI | TcI | [3] |
| SoCal3 | KR135421 | COII-NDI | TcI | [3] |
| Esc2 | KR135422 | COII-NDI | TcI | [3] |
| Esc26 | KR135424 | COII-NDI | TcI | [3] |
| Esc46 | KR135425 | COII-NDI | TcI | [3] |
| Colombia | EU302213 | COII-NDI | TcI | [2] |
| TEH cl2 cl92 | AF359009 | COII-NDI | TcI | [1] |
| OPS21 cl11 | AF359020 | COII-NDI | TcI | [1] |
| St5 | EU302209 | COII-NDI | TcI | [2] |
| CA R | GU212877 | COII-NDI | TcI | [4] |
| USA Armadillo | GU212880 | COII-NDI | TcI | [4] |
| USA Opossum | GU212879 | COII-NDI | TcI | [4] |
| JR cl4 | HQ604893 | COII-NDI | TcI | [5] |
| Tulahuen cl2 | AF359042 | COII-NDI | TcVI | [1] |
| CL F11F5 | AF359041 | COII-NDI | TcVI | [1] |
| VFRA1 cl1 | HQ604885 | COII-NDI | TcVI | [6] |
| 208 | EU302220 | COII-NDI | TcVI | [2] |
| Ab15 | EU302219 | COII-NDI | TcVI | [2] |
| Tul-K | EU302221 | COII-NDI | TcVI | [2] |
| CL Brener | DQ343645 | COII-NDI | TcVI | [7] |
| X110/8 | AF359034 | COII-NDI | TcIII | [1] |
| Ab3-10 | EU302218 | COII-NDI | TcII | [2] |
| EPP | AF359044 | COII-NDI | TcV | [1] |
| SO3 cl5 | AF359039 | COII-NDI | TcV | [1] |
| PSC-O | AF359040 | COII-NDI | TcV | [1] |
| BRJ | EU302217 | COII-NDI | TcIV | [2] |
| M6241 cl6 | AF359032 | COII-NDI | TcIII | [1] |
| CM 17 | AF359033 | COII-NDI | TcIII | [1] |
| EP 255 | AF359031 | COII-NDI | TcIV | [1] |
| 10R26 | HQ604907 | COII-NDI | TcIV | [5] |
| CANIII cl1 | AF359030 | COII-NDI | TcIV | [1] |
| Sjmc4 | HQ604902 | COII-NDI | TcIII | [5] |

| | | | | |
|---|---|---|---|---|
| Esc19 | KR135423 | COII-NDI | TcIV | [3] |
| Florida C16 | AF359010 | COII-NDI | TcIV | [1] |
| FH4 | GU212882 | COII-NDI | TcIV | [4] |
| USA Dog Y | GU212899 | COII-NDI | TcIV | [4] |
| 92122102R | HQ604909 | COII-NDI | TcIV | [5] |
| STC 10R | GU212900 | COII-NDI | TcIV | [4] |
| 93071502R | *GU212874 | COII-NDI | TcIV | [4] |
| GA Rac 134 | *GU212892 | COII-NDI | TcIV | [4] |
| GA Rac 143 | *GU212893 | COII-NDI | TcIV | [4] |
| GA Arm 20 | *GU212890 | COII-NDI | TcIV | [4] |
| STC 35R | *GU212897 | COII-NDI | TcIV | [4] |
| StC10R cl1 | *HQ604910 | COII-NDI | TcIV | [5] |
| FL Rac 15 | *GU212886 | COII-NDI | TcIV | [4] |
| 93040701R | *GU212872 | COII-NDI | TcIV | [4] |
| GA Rac 69 | *GU212895 | COII-NDI | TcIV | [4] |
| FL Rac 30 | *GU212887 | COII-NDI | TcIV | [4] |
| FL Rac 7 | *GU212888 | COII-NDI | TcIV | [4] |
| FL Rac 9 | *GU212889 | COII-NDI | TcIV | [4] |
| GA Rac 107 | *GU212891 | COII-NDI | TcIV | [4] |
| TN Rac 18 | *GU212898 | COII-NDI | TcIV | [4] |
| Dog Theis | *GU212881 | COII-NDI | TcIV | [4] |
| OK Dog | *GU212901 | COII-NDI | TcIV | [4] |
| Caesar Dog | *GU212894 | COII-NDI | TcIV | [4] |
| 92101601P | *GU212870 | COII-NDI | TcIV | [4] |
| FL Opo 2 | *GU212884 | COII-NDI | TcIV | [4] |
| 93070103P | *GU212873 | COII-NDI | TcIV | [4] |
| Samantha Dog | *GU212896 | COII-NDI | TcIV | [4] |
| FL Opo 3 | *GU212885 | COII-NDI | TcIV | [4] |
| EP 255 | AF358943 | DHFR-TS | TcIV | [1] |
| CANIII cl1 | AF358941 | DHFR-TS | TcIV | [1] |

| | | | | |
|---|---|---|---|---|
| GA Rac 143 | GU212915 | DHFR-TS | TcIV | [4] |
| 93072805R | GU212907 | DHFR-TS | TcIV | [4] |
| GA Rac 69 | GU212917 | DHFR-TS | TcIV | [4] |
| FL Rac 13 | GU212910 | DHFR-TS | TcIV | [4] |
| FL Rac 30 | GU212911 | DHFR-TS | TcIV | [4] |
| STC 35R | GU212919 | DHFR-TS | TcIV | [4] |
| 92122102R | GU212903 | DHFR-TS | TcIV | [4] |
| 93040701R | GU212904 | DHFR-TS | TcIV | [4] |
| 93071502R | GU212905 | DHFR-TS | TcIV | [4] |
| Caesar Dog | GU212932 | DHFR-TS | TcIV | [4] |
| TN Rac 18 | GU212920 | DHFR-TS | TcIV | [4] |
| FL Rac 7 | GU212912 | DHFR-TS | TcIV | [4] |
| FL Rac 9 | GU212913 | DHFR-TS | TcIV | [4] |
| GA Rac 45 | GU212916 | DHFR-TS | TcIV | [4] |
| YuYu clone 1 | AY785655 | DHFR-TS | TcI | Not published |
| CM 17 | AF358946 | DHFR-TS | TcIII | [1] |
| X110/8 | AF358947 | DHFR-TS | TcIII | [1] |
| EPP | AF358965 | DHFR-TS | TcV | [1] |
| SO3 cl5 | AF358957 | DHFR-TS | TcV | [1] |
| M5631 cl5 | AF358945 | DHFR-TS | TcIII | [1] |
| PSC-O | AF358959 | DHFR-TS | TcV | [1] |
| M6241 cl6 | AF358944 | DHFR-TS | TcIII | [1] |
| Y82R | JN942603 | DHFR-TS | TcII | [8] |
| TEH cl2 cl92 | AF358926 | DHFR-TS | TcI | [1] |
| OPS21 cl11 | AF358932 | DHFR-TS | TcI | [1] |
| CUICA cl1 | AF358936 | DHFR-TS | TcI | [1] |
| Florida C1F8 | GU212926 | DHFR-TS | TcI | [4] |
| 93070103P | GU212906 | DHFR-TS | TcI | [4] |
| X10 cl1 | AF358929 | DHFR-TS | TcI | [1] |
| 133 79 cl7 | AF358935 | DHFR-TS | TcI | [1] |
| SC13 | AF358942 | DHFR-TS | TcI | [1] |
| CL Brener | XM_814525 | DHFR-TS | TcVI | [9] |
| YUSPR | JN942599 | DHFR-TS | TcII | [8] |
| YUSP | JN942598 | DHFR-TS | TcII | [8] |
| Y | EU302241 | DHFR-TS | TcII | [2] |
| Tul L | EU302238 | DHFR-TS | TcII | [2] |
| CL F11F5 | AF358961 | DHFR-TS | TcVI | [1] |
| SC43 cl1 clone 1 | AY785645 | DHFR-TS | TcV | unpublished |
| MSC2 | AF358954 | DHFR-TS | TcII | [1] |
| Esmeraldo cl3 | AF358948 | DHFR-TS | TcII | [1] |
| *Trypanosoma cruzi marinkellei* 593 B3 | AF358966 | DHFR-TS | outgroup | [1] |
| 167b | AY092825 | MSH2 | TcII | [10] |
| 115a | AY092835 | MSH2 | TcII | [10] |
| GA Sk 1 | GU213035 | MSH2 | TcIV | [4] |
| Texas Theis | GU213031 | MSH2 | TcIV | [4] |
| FL Rac 5 | GU213014 | MSH2 | TcIV | [4] |
| STC 35R | GU213024 | MSH2 | TcIV | [4] |
| Maryland Rac | GU213022 | MSH2 | TcIV | [4] |
| Caesar Dog | GU212989 | MSH2 | TcIV | [4] |
| Dog Theis | GU212990 | MSH2 | TcIV | [4] |
| 93072805R | GU213009 | MSH2 | TcIV | [4] |
| 93071502R | GU213008 | MSH2 | TcIV | [4] |

| | | | | |
|---|---|---|---|---|
| 92122102R | GU213005 | MSH2 | TcIV | [4] |
| GA Rac 69 | GU213021 | MSH2 | TcIV | [4] |
| 93040701R | GU213006 | MSH2 | TcIV | [4] |
| Smokey | GU212994 | MSH2 | TcIV | [4] |
| OK Dog | GU212992 | MSH2 | TcIV | [4] |
| STC 10R | GU213023 | MSH2 | TcIV | [4] |
| USA Armadillo | GU213034 | MSH2 | TcI | [4] |
| Sylvio X10/cl1 | CP015651 | MSH2 | TcI | [11] |
| colombiana | AY092836 | MSH2 | TcI | [10] |
| Armadillo 1973 | GU213032 | MSH2 | TcI | [4] |
| 93053103R | GU213007 | MSH2 | TcI | [4] |
| M5631 cl5 | AY540734 | MSH2 | TcIII | [12] |
| P255 | DQ021899 | MSH2 | TcV | [12] |
| 231 | AY092826 | MSH2 | TcV/VI | [10] |
| CL Brener | XM_806693 | MSH2 | TcVI | [9] |
| Tulahuen | AY005798 | MSH2 | TcV | [13] |
| MN cl.2 | DQ021898 | MSH2 | TcV | [12] |
| MN cl2 | AY540736 | MSH2 | TcV | [12] |
| 115b | AY092828 | MSH2 | TcV/VI | [10] |
| CanIII cl1 | AY540738 | MSH2 | TcIV | [12] |
| M4167 | AY540733 | MSH2 | TcIV | [12] |
| TU18 cl2 | AY540731 | MSH2 | TcIII/V/VI | [12] |
| 593 | AY092830 | MSH2 | TcII | [10] |
| 577 | AY092831 | MSH2 | TcII | [10] |
| CBB cl3 | AY540740 | MSH2 | TcII | [12] |
| 239b | AY092837 | MSH2 | TcII | [10] |
| 239a | AY092838 | MSH2 | TcII | [10] |
| Y | AY540737 | MSH2 | TcII | [12] |
| SC43 cl1 | AY540735 | MSH2 | TcV | [12] |
| S034_cl4_B | HQ859547 | 6529.310 | TcI | [14] |
| 133_79_cl7_B | HQ859545 | 6529.310 | TcI | [14] |
| SC13 | HQ859550 | 6529.310 | TcI | [14] |
| 133_79_cl7_A | HQ859544 | 6529.310 | TcI | [14] |
| S034_cl4_A | HQ859546 | 6529.310 | TcI | [14] |
| 39_EPP_A | HQ859561 | 6529.310 | TcV | [14] |
| M6241_cl6 | HQ859553 | 6529.310 | TcIII | [14] |
| CL Brener | XM_803199 | 6529.310 | TcVI | [9] |
| M5631 cl5 | HQ859554 | 6529.310 | TcIII | [14] |
| CANIII_cl1 | HQ859551 | 6529.310 | TcIV | [14] |
| EP_255 | HQ859552 | 6529.310 | TcIV | [14] |
| TCC 1994 | KT327245 | 6529.310 | TcBAT | [15] |
| ESMERALDO_cl3_A | HQ859555 | 6529.310 | TcII | [14] |
| TU18_cl2_B | HQ859560 | 6529.310 | TcII/V/VI | [14] |
| EPP_B | HQ859562 | 6529.310 | TcV | [14] |
| CL_Brener_A | HQ859565 | 6529.310 | TcVI | [14] |
| ESMERALDO_cl3_B | HQ859556 | 6529.310 | TcII | [14] |
| CBB_cl3_B | HQ859558 | 6529.310 | TcII | [14] |
| TU18_cl2_A | HQ859559 | 6529.310 | TcII/V/VI | [14] |

| | | | | |
|---|---|---|---|---|
| *Trypanosoma cruzi marinkellei* TCC 344 | KT327246 | 6529.310 | Outgroup | [15] |

| **Mitochondrial loci** (COII-NDI) | | | | | | | | |
|---|---|---|---|---|---|---|---|---|
| | *T.cruzi* | TcII | TcIII-IV | TcI North Am. | TcI South Am. | TcI | TcIV-USA | Posterior likelihood |
| Strict | 3.49 (2.09-4.8) | 0.209 (0.07-0.36) | 0.542 (0.28-0.83) | 0.263 (0.11-0.41) | 0.405 (0.21-0.61) | 0.475 (0.26-0.71) | 0.12 (0.03-0.21 | -4659.033 |
| Relaxed | 3.52 (1.36-5.8) | 0.244 (0.047-0.51) | 0.56 (0.17-1.03) | 0.315 (0.086-0.59) | 0.454 (0.13-0.81) | 0.59 (0.19-1.07) | 0.159 (0.032-0.34) | -4656.375 |

| **NUCLEAR (**3 loci) | | | | | | |
|---|---|---|---|---|---|---|
| | *T.cruzi* | TcII | TcIII-IV (not informative) | TcI | TcIV-USA | Posterior likelihood |
| Strict | 1.63 (0.95-2.33) | 0.43 (0.19-0.69) | 1.15 (0.67-1.7) | 0.073 (0.001-0.18) | 0.11 (0.02-0.22) | -5124.638 |

**S3 Table. Bayesian estimates of divergence (time in millions of years) for the main *T. cruzi* lineages and TcIV-USA.**

Nuclear data consisted of 2,757 concatenated nucleotides, whereas mitochondrial data set consisted of 1,161 nucleotides. Due to inconsistency of amplifying all 3 nuclear loci in each sample, for the nuclear analyses we only included isolates that had all three nuclear loci. Both analyses were calibrated with the divergence estimate of 6.23 million years ago (with 1.5 million years of standard deviation with a prior normal distribution for the tmrca) between *T. cruzi* and *T. c. marinkellei* that was estimated from our previous analysis[21]. Parameters: HKY as DNA substitution model, gamma with invariant sites as the site heterogeneity model, with 4 gamma categories, data was partitioned into codons. Convergence could not be attained for the nuclear relaxed analysis thus data is not shown. TcI North Am: clade of TcI that is exclusively composed of samples from North America. TcI South Am: clade of TcI that is exclusively composed of samples from South America. Tex 16 was not included in the analysis due to conflicting phylogenies between nuclear trees (Technical Supplement Figures 1-3).

Fig S1

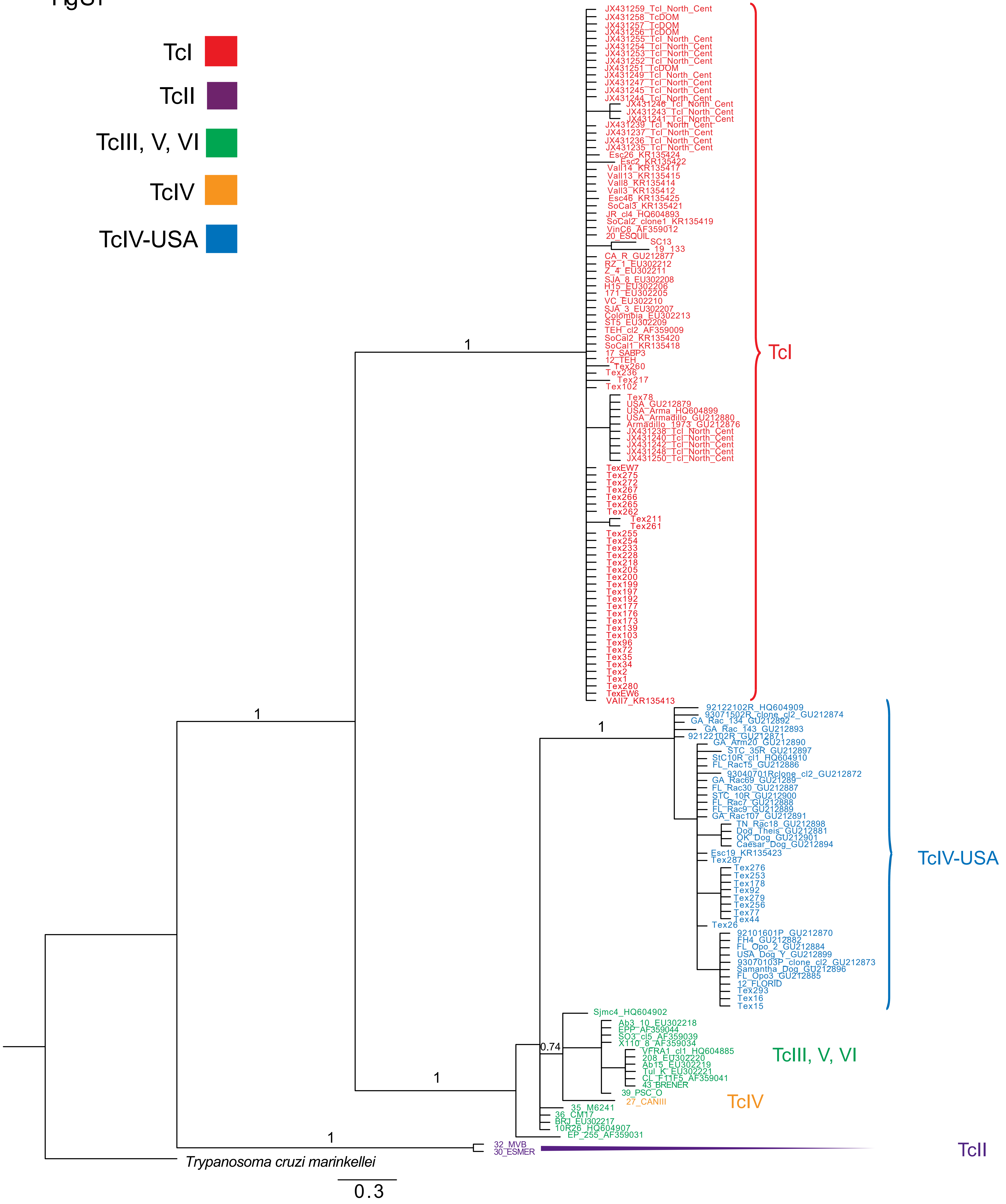

TcI
TcII
TcIII, V, VI
TcIV
TcIV-USA
JX431259_TcI_North_Cent
JX431258_TcDOM
JX431257_TcDOM
JX431256_TcDOM
JX431255_TcI_North_Cent
JX431254_TcI_North_Cent
JX431253_TcI_North_Cent
JX431252_TcI_North_Cent
JX431251_TcDOM
JX431249_TcI_North_Cent
JX431247_TcI_North_Cent
JX431245_TcI_North_Cent
JX431244_TcI_North_Cent
JX431246_TcI_North_Cent
JX431243_TcI_North_Cent
JX431241_TcI_North_Cent
JX431239_TcI_North_Cent
JX431237_TcI_North_Cent
JX431236_TcI_North_Cent
JX431235_TcI_North_Cent
Esc26_KR135424
Esc2_KR135422
Vall14_KR135417
Vall13_KR135415
Vall8_KR135414
Vall3_KR135412
Esc46_KR135425
SoCal3_KR135421
JR_cl4_HQ604893
SoCal2_clone1_KR135419
VinC6_AF359012
20_ESQUIL
SC13
19_133
CA_R_GU212877
RZ_1_EU302212
Z_4_EU302211
SJA_8_EU302208
H15_EU302206
171_EU302205
VC_EU302210
SJA_3_EU302207
Colombia_EU302213
ST5_EU302209
TEH_cl2_AF359009
SoCal2_KR135420
SoCal1_KR135418
17_SABP3
12_TEH
Tex260
Tex236
Tex217
Tex102
Tex78
USA_GU212879
USA_Arma_HQ604899
USA_Armadillo_GU212880
Armadillo_1973_GU212876
JX431238_TcI_North_Cent
JX431240_TcI_North_Cent
JX431242_TcI_North_Cent
JX431248_TcI_North_Cent
JX431250_TcI_North_Cent
TexEW7
Tex275
Tex272
Tex267
Tex266
Tex265
Tex262
Tex211
Tex261
Tex255
Tex254
Tex233
Tex228
Tex218
Tex205
Tex200
Tex199
Tex197
Tex192
Tex177
Tex176
Tex173
Tex139
Tex103
Tex96
Tex72
Tex35
Tex34
Tex2
Tex1
Tex280
TexEW6
VAll7_KR135413
TcI
1
1
1
92122102R_HQ604909
93071502R_clone_cl2_GU212874
GA_Rac_134_GU212892
GA_Rac_143_GU212893
92122102R_GU212871
GA_Arm20_GU212890
STC_35R_GU212897
StC10R_cl1_HQ604910
FL_Rac15_GU212886
93040701Rclone_cl2_GU212872
GA_Rac69_GU212895
FL_Rac30_GU212887
STC_10R_GU212900
FL_Rac7_GU212888
FL_Rac9_GU212889
GA_Rac107_GU212891
TN_Rac18_GU212898
Dog_Theis_GU212881
OK_Dog_GU212901
Caesar_Dog_GU212894
Esc19_KR135423
Tex287
Tex276
Tex253
Tex178
Tex92
Tex279
Tex256
Tex77
Tex44
Tex26
92101601P_GU212870
FH4_GU212882
FL_Opo_2_GU212884
USA_Dog_Y_GU212899
93070103P_clone_cl2_GU212873
Samantha_Dog_GU212896
FL_Opo3_GU212885
12_FLORID
Tex293
Tex16
Tex15
TcIV-USA
Sjmc4_HQ604902
Ab3_10_EU302218
EPP_AF359044
SO3_cl5_AF359039
X110_8_AF359034
VFRA1_cl1_HQ604885
208_EU302220
Ab15_EU302219
Tul_K_EU302221
CL_F11F5_AF359041
43_BRENER
39_PSC_O
27_CANIII
35_M6241
36_CM17
BRJ_EU302217
10R26_HQ604907
EP_255_AF359031
0.74
1
TcIII, V, VI
TcIV
1
32_MVB
30_ESMER
TcII
*Trypanosoma cruzi marinkellei*
0.3

Fig S2

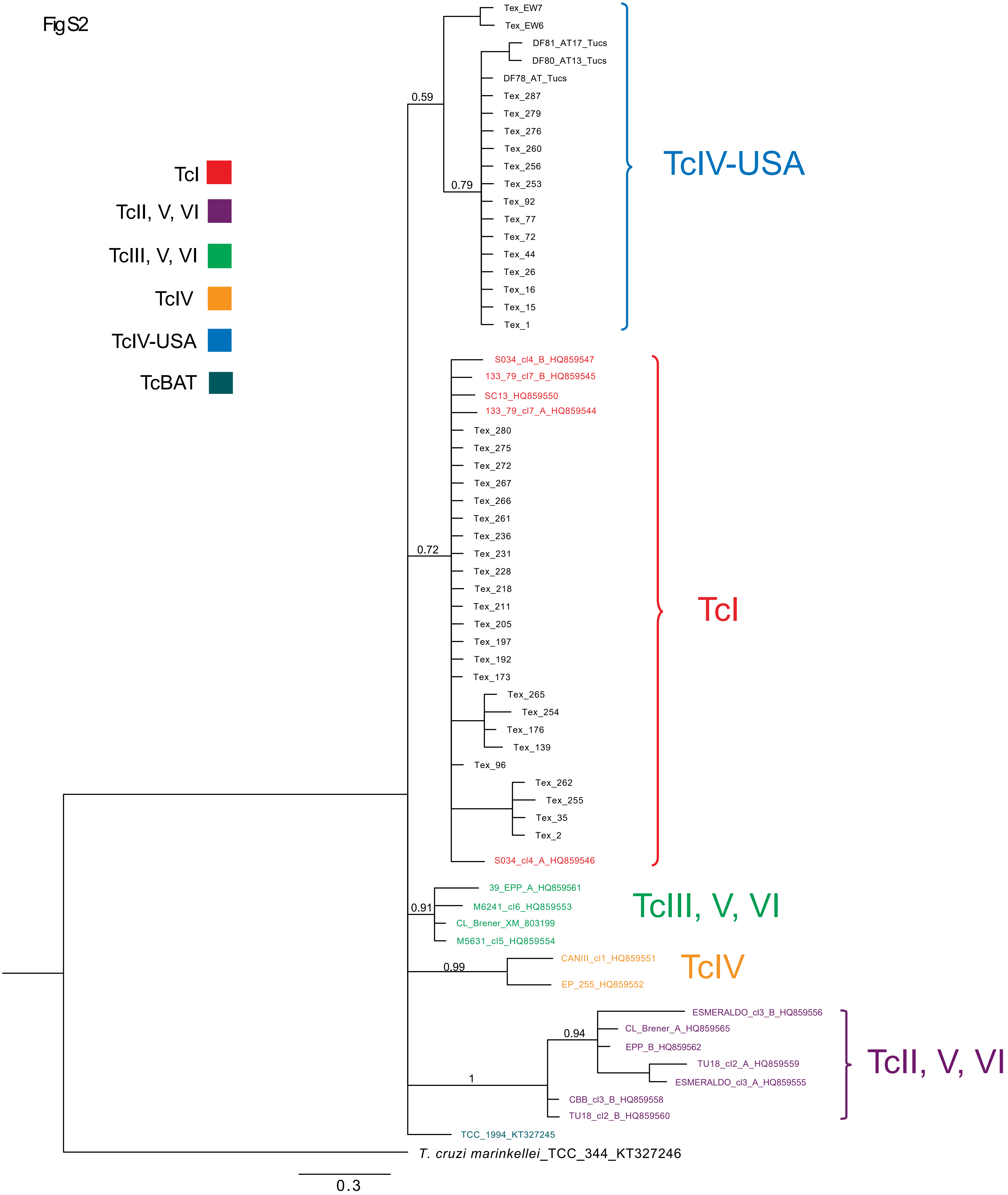

TcI
TcII, V, VI
TcIII, V, VI
TcIV
TcIV-USA
TcBAT
Tex_EW7
Tex_EW6
DF81_AT17_Tucs
DF80_AT13_Tucs
DF78_AT_Tucs
Tex_287
Tex_279
Tex_276
Tex_260
Tex_256
Tex_253
Tex_92
Tex_77
Tex_72
Tex_44
Tex_26
Tex_16
Tex_15
Tex_1
0.59
0.79
TcIV-USA
S034_cl4_B_HQ859547
133_79_cl7_B_HQ859545
SC13_HQ859550
133_79_cl7_A_HQ859544
Tex_280
Tex_275
Tex_272
Tex_267
Tex_266
Tex_261
Tex_236
Tex_231
Tex_228
Tex_218
Tex_211
Tex_205
Tex_197
Tex_192
Tex_173
Tex_265
Tex_254
Tex_176
Tex_139
Tex_96
Tex_262
Tex_255
Tex_35
Tex_2
S034_cl4_A_HQ859546
0.72
TcI
39_EPP_A_HQ859561
M6241_cl6_HQ859553
CL_Brener_XM_803199
M5631_cl5_HQ859554
0.91
TcIII, V, VI
CANIII_cl1_HQ859551
EP_255_HQ859552
0.99
TcIV
ESMERALDO_cl3_B_HQ859556
CL_Brener_A_HQ859565
EPP_B_HQ859562
TU18_cl2_A_HQ859559
ESMERALDO_cl3_A_HQ859555
CBB_cl3_B_HQ859558
TU18_cl2_B_HQ859560
0.94
1
TcII, V, VI
TCC_1994_KT327245
T. cruzi marinkellei_TCC_344_KT327246
0.3

Fig S3

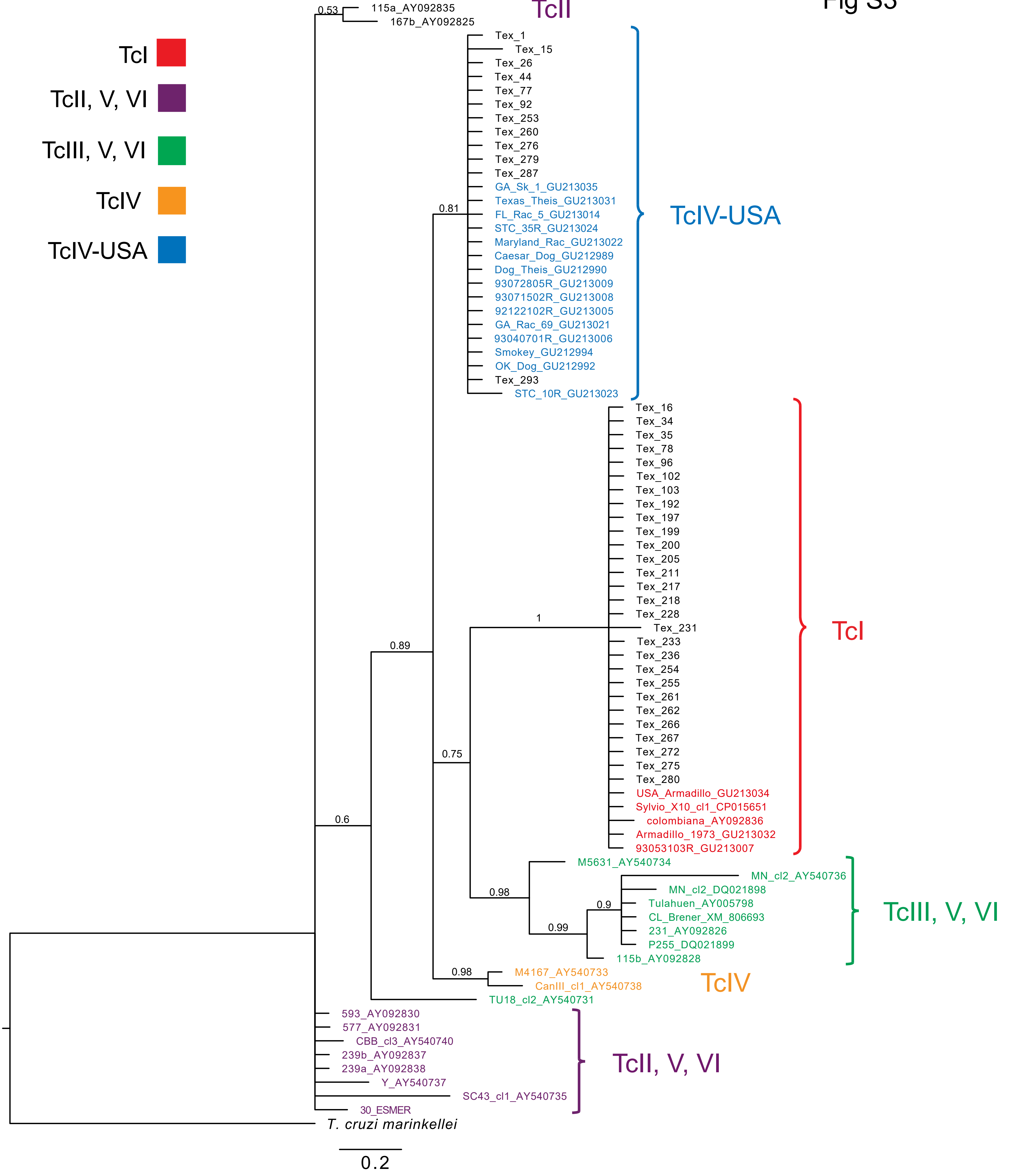
TcI
TcII, V, VI
TcIII, V, VI
TcIV
TcIV-USA
TcII
0.53
115a_AY092835
167b_AY092825
Tex_1
Tex_15
Tex_26
Tex_44
Tex_77
Tex_92
Tex_253
Tex_260
Tex_276
Tex_279
Tex_287
GA_Sk_1_GU213035
Texas_Theis_GU213031
0.81
FL_Rac_5_GU213014
STC_35R_GU213024
Maryland_Rac_GU213022
Caesar_Dog_GU212989
Dog_Theis_GU212990
93072805R_GU213009
93071502R_GU213008
92122102R_GU213005
GA_Rac_69_GU213021
93040701R_GU213006
Smokey_GU212994
OK_Dog_GU212992
Tex_293
STC_10R_GU213023
TcIV-USA
Tex_16
Tex_34
Tex_35
Tex_78
Tex_96
Tex_102
Tex_103
Tex_192
Tex_197
Tex_199
Tex_200
Tex_205
Tex_211
Tex_217
Tex_218
Tex_228
1
Tex_231
0.89
Tex_233
Tex_236
Tex_254
Tex_255
Tex_261
Tex_262
Tex_266
Tex_267
0.75
Tex_272
Tex_275
Tex_280
USA_Armadillo_GU213034
0.6
Sylvio_X10_cl1_CP015651
colombiana_AY092836
Armadillo_1973_GU213032
93053103R_GU213007
TcI
M5631_AY540734
MN_cl2_AY540736
MN_cl2_DQ021898
0.98
0.9
Tulahuen_AY005798
CL_Brener_XM_806693
0.99
231_AY092826
P255_DQ021899
115b_AY092828
TcIII, V, VI
0.98
M4167_AY540733
CanIII_cl1_AY540738
TcIV
TU18_cl2_AY540731
593_AY092830
577_AY092831
CBB_cl3_AY540740
239b_AY092837
239a_AY092838
Y_AY540737
SC43_cl1_AY540735
30_ESMER
TcII, V, VI
T. cruzi marinkellei
0.2

Fig S4

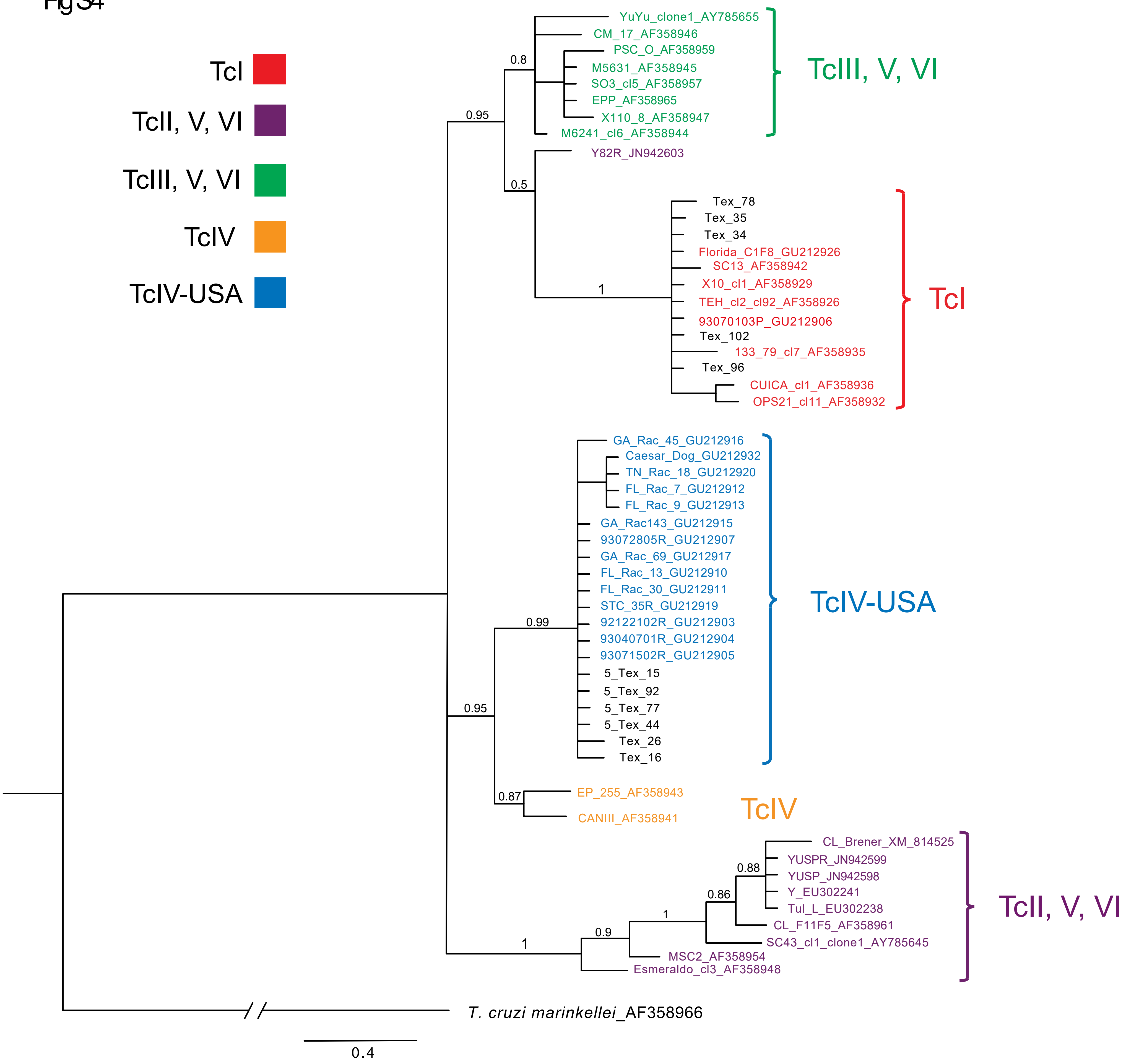

TcI
TcII, V, VI
TcIII, V, VI
TcIV
TcIV-USA
YuYu_clone1_AY785655
CM_17_AF358946
PSC_O_AF358959
M5631_AF358945
SO3_cl5_AF358957
EPP_AF358965
X110_8_AF358947
M6241_cl6_AF358944
Y82R_JN942603
Tex_78
Tex_35
Tex_34
Florida_C1F8_GU212926
SC13_AF358942
X10_cl1_AF358929
TEH_cl2_cl92_AF358926
93070103P_GU212906
Tex_102
133_79_cl7_AF358935
Tex_96
CUICA_cl1_AF358936
OPS21_cl11_AF358932
GA_Rac_45_GU212916
Caesar_Dog_GU212932
TN_Rac_18_GU212920
FL_Rac_7_GU212912
FL_Rac_9_GU212913
GA_Rac143_GU212915
93072805R_GU212907
GA_Rac_69_GU212917
FL_Rac_13_GU212910
FL_Rac_30_GU212911
STC_35R_GU212919
92122102R_GU212903
93040701R_GU212904
93071502R_GU212905
5_Tex_15
5_Tex_92
5_Tex_77
5_Tex_44
Tex_26
Tex_16
EP_255_AF358943
CANIII_AF358941
CL_Brener_XM_814525
YUSPR_JN942599
YUSP_JN942598
Y_EU302241
Tul_L_EU302238
CL_F11F5_AF358961
SC43_cl1_clone1_AY785645
MSC2_AF358954
Esmeraldo_cl3_AF358948
T. cruzi marinkellei_AF358966
0.8
0.95
0.5
1
0.99
0.95
0.87
0.88
0.86
1
0.9
1
0.4

Fig S5

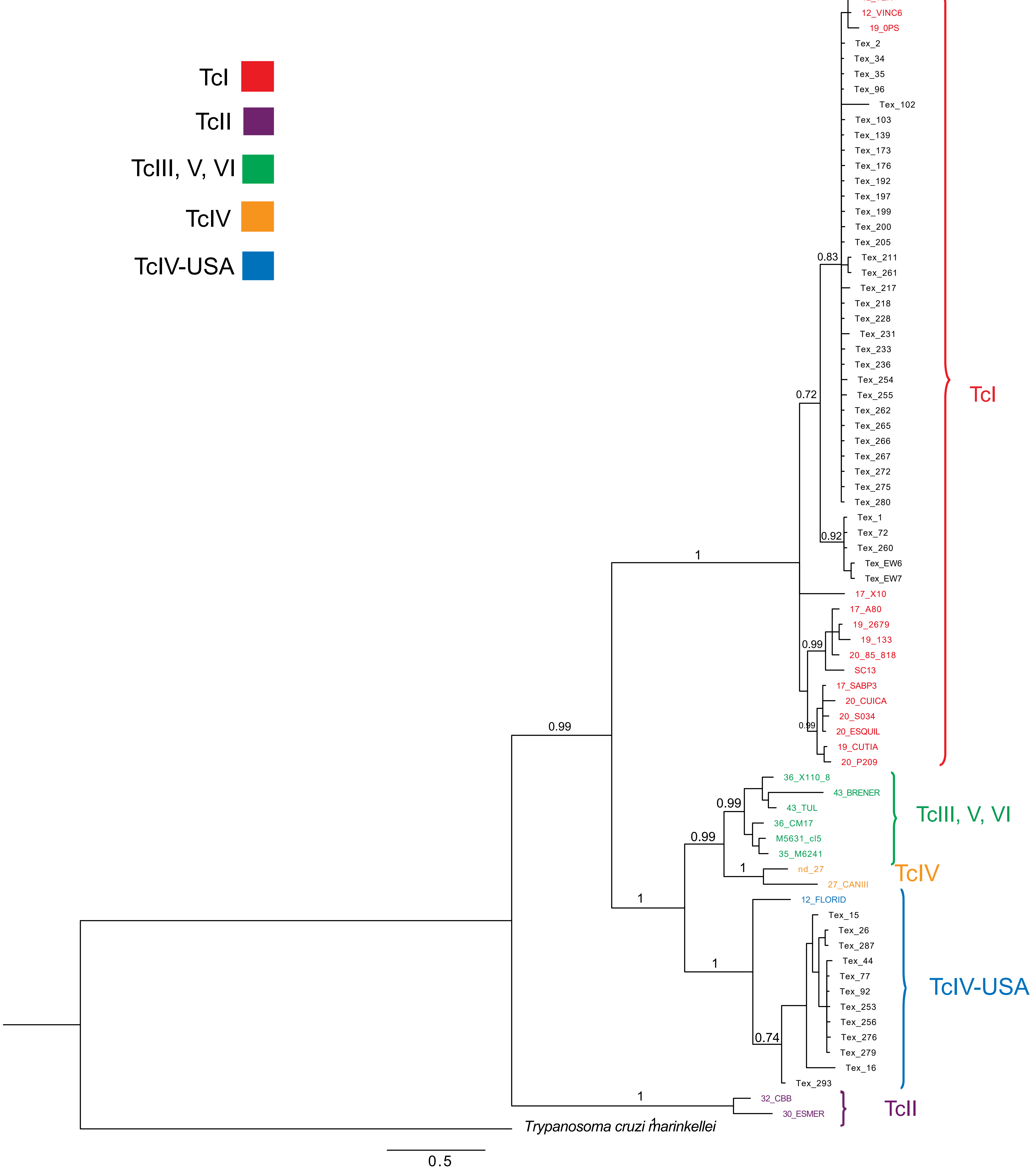
TcI
TcII
TcIII, V, VI
TcIV
TcIV-USA
12_TEH
12_VINC6
19_0PS
Tex_2
Tex_34
Tex_35
Tex_96
Tex_102
Tex_103
Tex_139
Tex_173
Tex_176
Tex_192
Tex_197
Tex_199
Tex_200
Tex_205
0.83
Tex_211
Tex_261
Tex_217
Tex_218
Tex_228
Tex_231
Tex_233
Tex_236
Tex_254
0.72
Tex_255
Tex_262
Tex_265
Tex_266
Tex_267
Tex_272
Tex_275
Tex_280
Tex_1
0.92
Tex_72
Tex_260
1
Tex_EW6
Tex_EW7
17_X10
17_A80
19_2679
19_133
0.99
20_85_818
SC13
17_SABP3
20_CUICA
20_S034
0.99
0.99
20_ESQUIL
19_CUTIA
20_P209
TcI
36_X110_8
43_BRENER
0.99
43_TUL
36_CM17
0.99
M5631_cl5
35_M6241
1
nd_27
27_CANIII
TcIII, V, VI
TcIV
1
12_FLORID
Tex_15
Tex_26
Tex_287
Tex_44
Tex_77
1
Tex_92
Tex_253
Tex_256
Tex_276
0.74
Tex_279
Tex_16
Tex_293
TcIV-USA
1
32_CBB
30_ESMER
TcII
Trypanosoma cruzi marinkellei
1
0.5

Fig S6

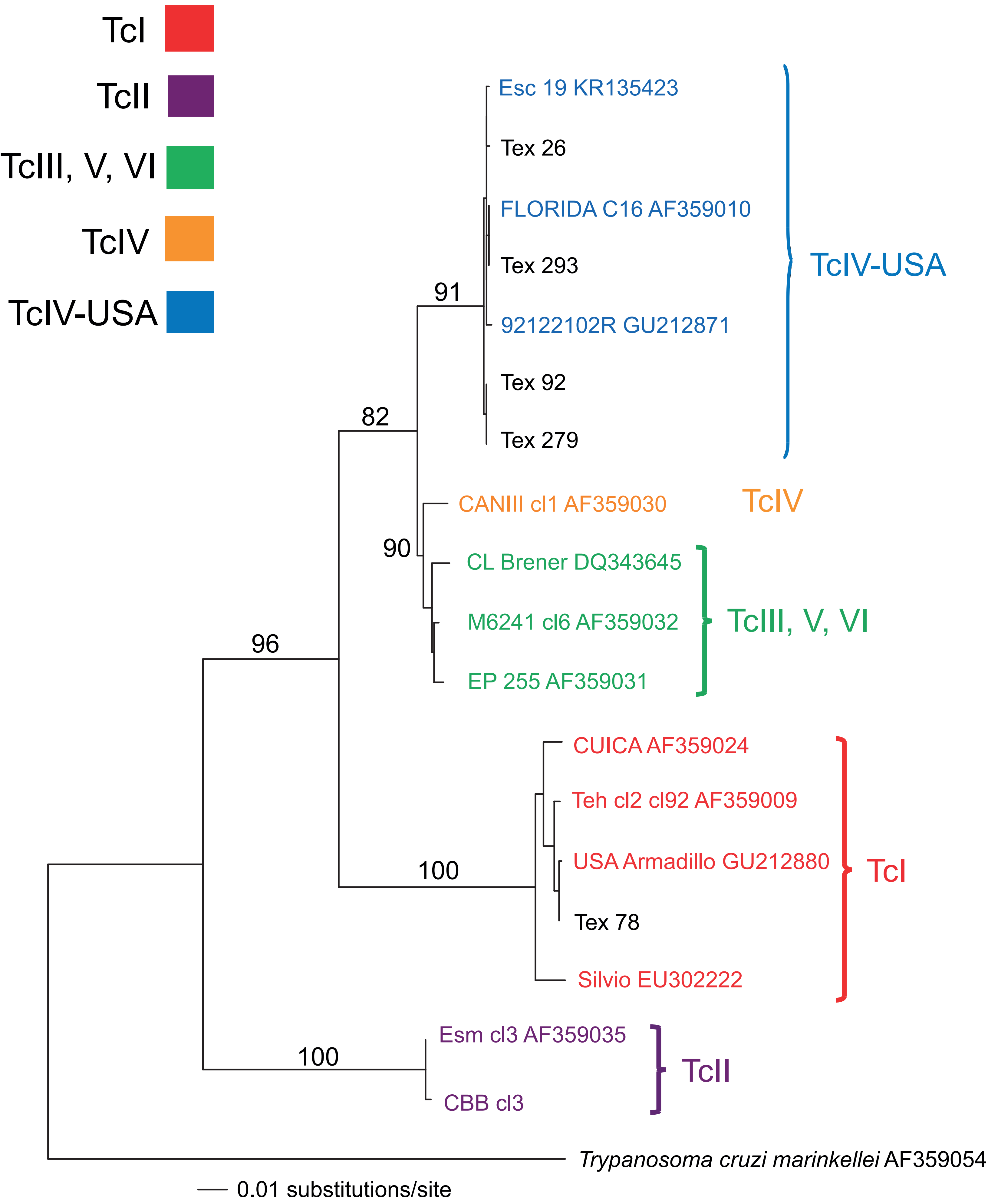
TcI
TcII
TcIII, V, VI
TcIV
TcIV-USA
Esc 19 KR135423
Tex 26
FLORIDA C16 AF359010
Tex 293
92122102R GU212871
Tex 92
Tex 279
TcIV-USA
91
82
CANIII cl1 AF359030
TcIV
90
CL Brener DQ343645
M6241 cl6 AF359032
EP 255 AF359031
TcIII, V, VI
96
CUICA AF359024
Teh cl2 cl92 AF359009
USA Armadillo GU212880
Tex 78
Silvio EU302222
TcI
100
Esm cl3 AF359035
CBB cl3
TcII
100
*Trypanosoma cruzi marinkellei* AF359054
0.01 substitutions/site

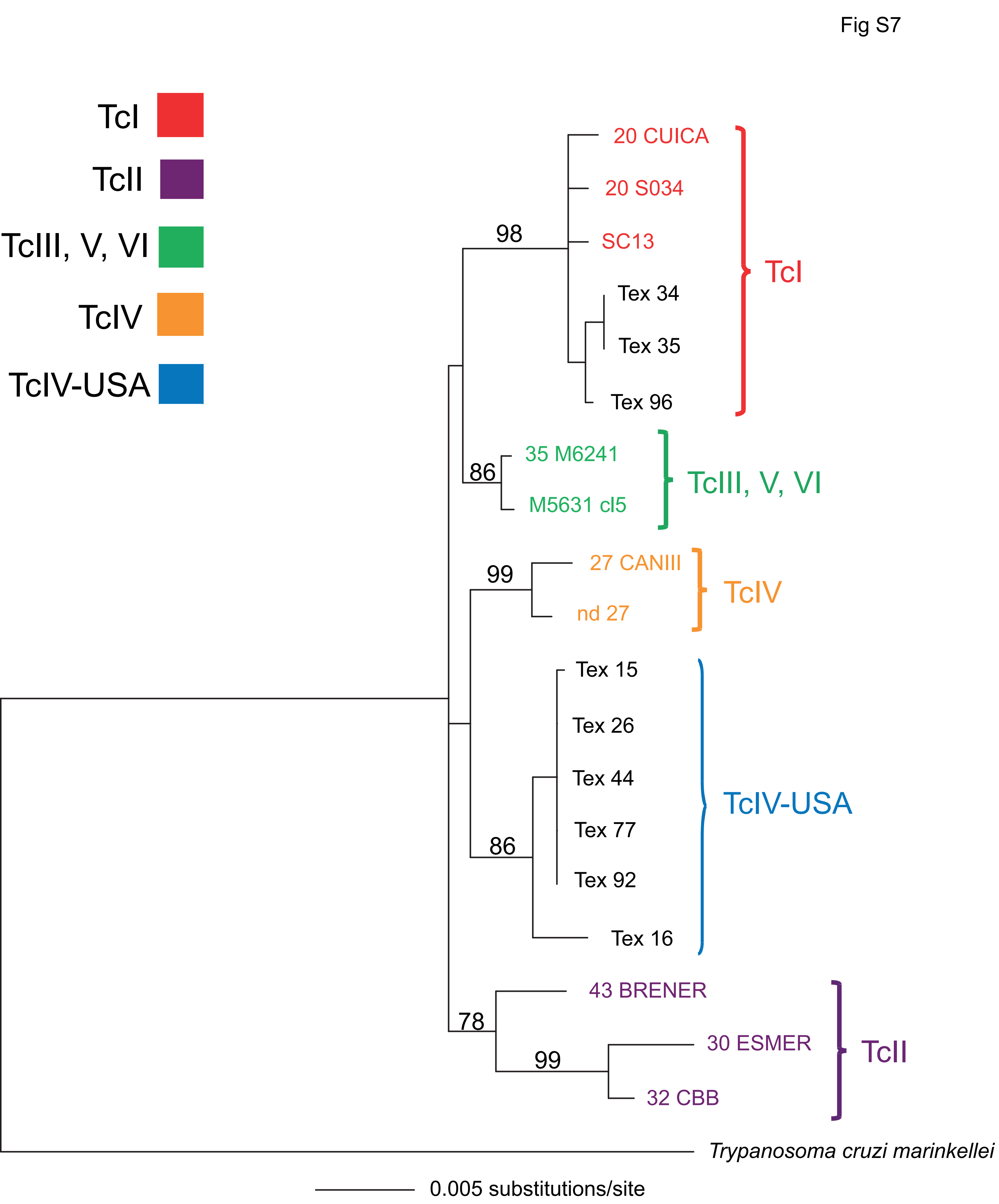
Fig S7
TcI
TcII
TcIII, V, VI
TcIV
TcIV-USA
20 CUICA
20 S034
SC13
Tex 34
Tex 35
Tex 96
98
35 M6241
M5631 cl5
86
27 CANIII
nd 27
99
Tex 15
Tex 26
Tex 44
Tex 77
Tex 92
Tex 16
86
43 BRENER
30 ESMER
32 CBB
78
99
Trypanosoma cruzi marinkellei
0.005 substitutions/site

Fig S8

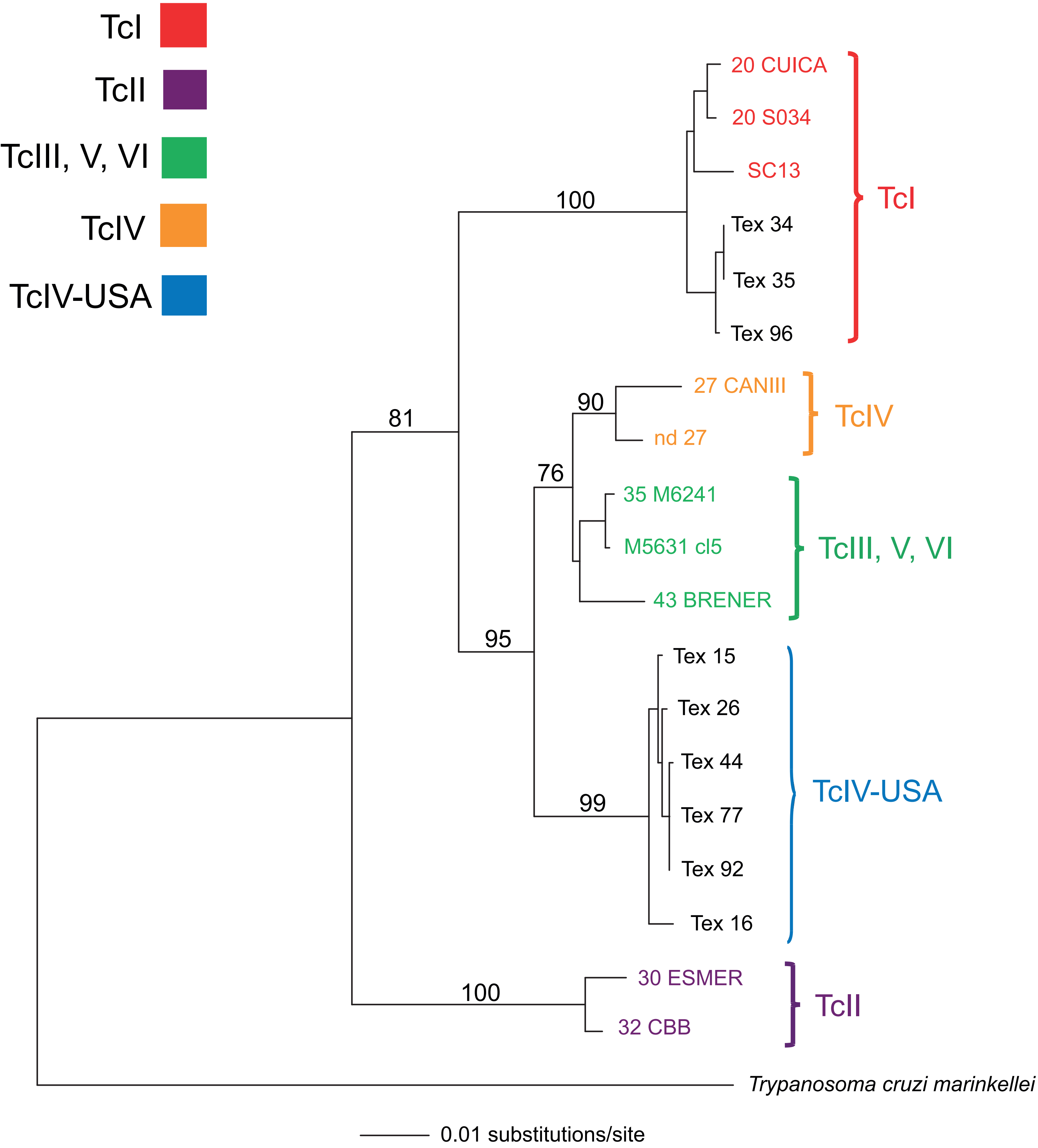
TcI
TcII
TcIII, V, VI
TcIV
TcIV-USA
100
20 CUICA
20 S034
SC13
Tex 34
Tex 35
Tex 96
TcI
81
90
27 CANIII
nd 27
TcIV
76
35 M6241
M5631 cl5
43 BRENER
TcIII, V, VI
95
Tex 15
Tex 26
Tex 44
Tex 77
Tex 92
Tex 16
99
TcIV-USA
100
30 ESMER
32 CBB
TcII
Trypanosoma cruzi marinkellei
0.01 substitutions/site

Fig S9

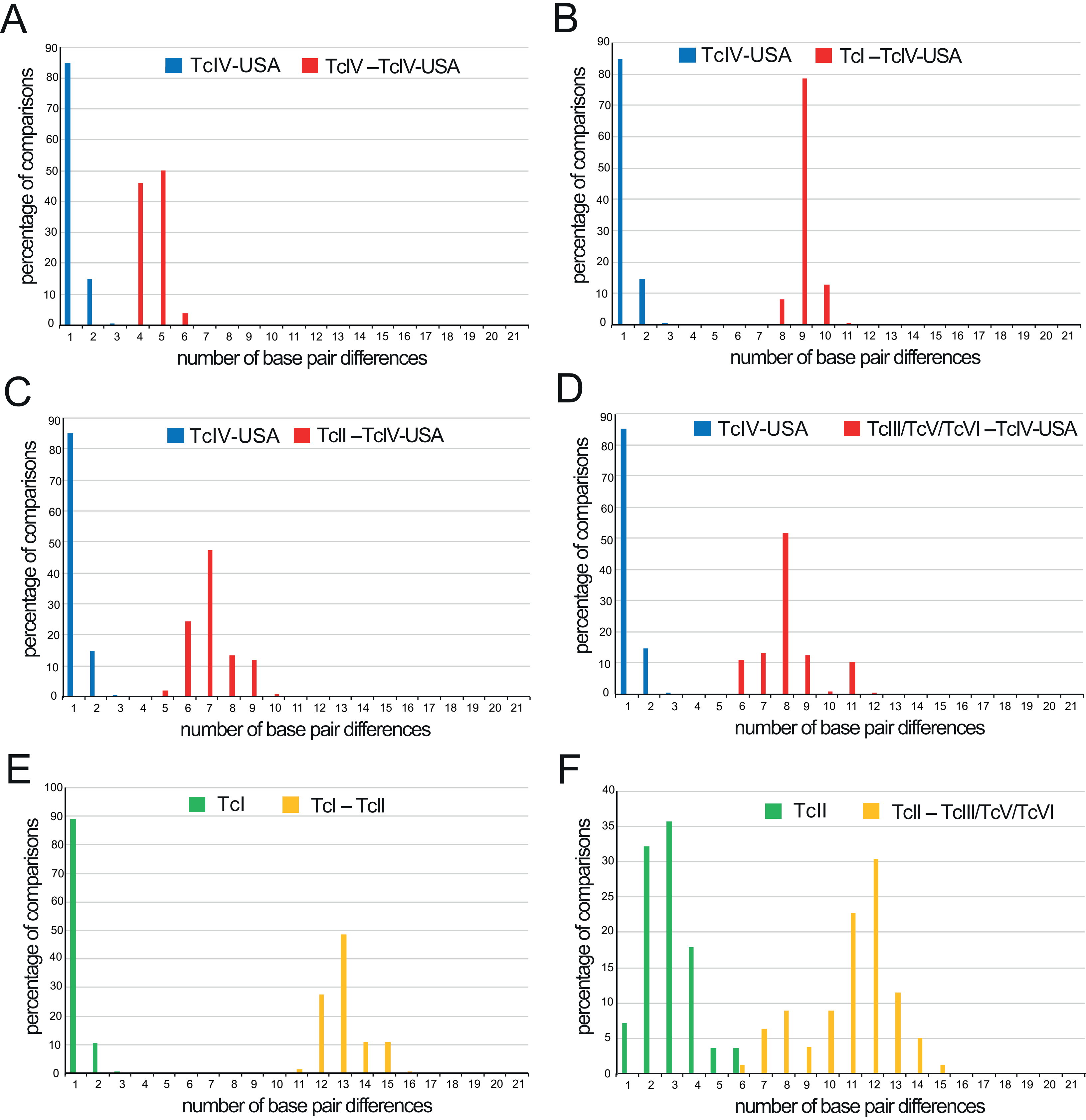
A
TcIV-USA
TcIV –TcIV-USA
percentage of comparisons
number of base pair differences
B
TcIV-USA
TcI –TcIV-USA
C
TcIV-USA
TcII –TcIV-USA
D
TcIV-USA
TcIII/TcV/TcVI –TcIV-USA
E
TcI
TcI – TcII
F
TcII
TcII – TcIII/TcV/TcVI

Fig S10

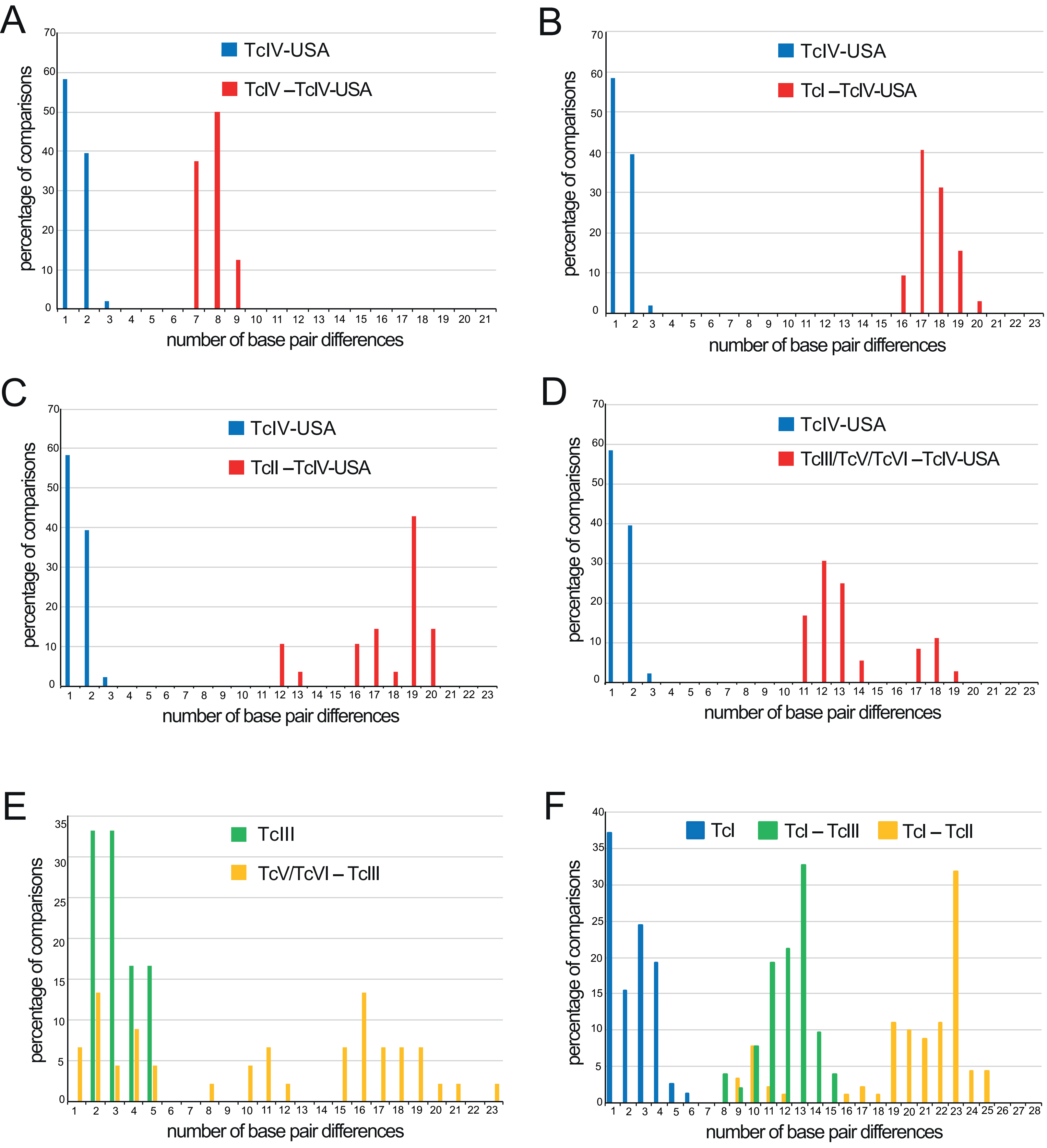

A
TcIV-USA
TcIV –TcIV-USA
percentage of comparisons
number of base pair differences
B
TcIV-USA
TcI –TcIV-USA
percentage of comparisons
number of base pair differences
C
TcIV-USA
TcII –TcIV-USA
percentage of comparisons
number of base pair differences
D
TcIV-USA
TcIII/TcV/TcVI –TcIV-USA
percentage of comparisons
number of base pair differences
E
TcIII
TcV/TcVI – TcIII
percentage of comparisons
number of base pair differences
F
TcI
TcI – TcIII
TcI – TcII
percentage of comparisons
number of base pair differences

Fig. S11

Potential Hybrid
Id: Texas 72
Collected 25 May 2011 in Medina county Texas, peridomestic/mixed Oak
Collected by Blake Sissel & Ed Wozniak, USA
Host: *Triatoma lecticularia*
MSH2

Position 165

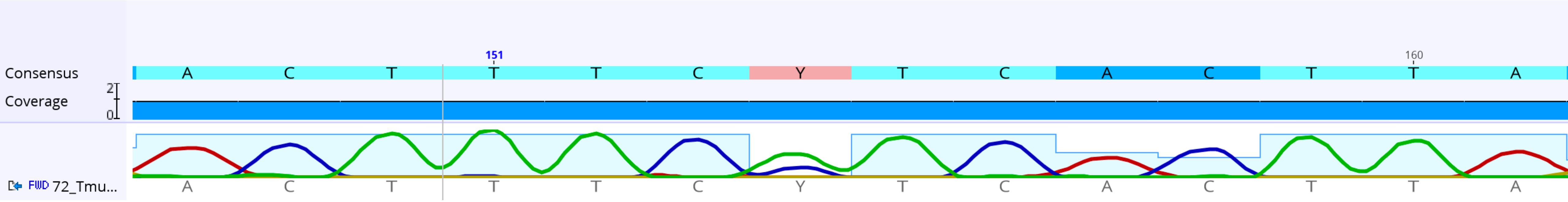


T: is from TcIV-USA allele
C: is from TcI allele

Potential Hybrid
Id: Texas 72
Collected 25 May 2011 in Medina county Texas, peridomestic/mixed Oak
Collected by Blake Sissel & Ed Wozniak, USA
Host: *Triatoma lecticularia*
MSH2

Position 288

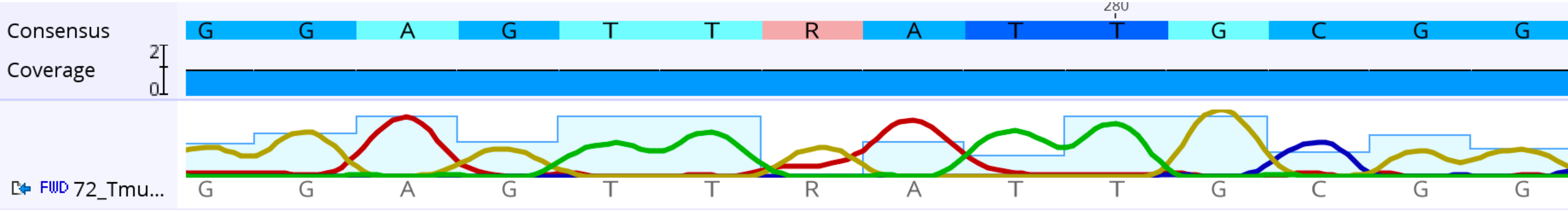


G: is from TcIV-USA allele
A: is from TcI allele

Potential Hybrid
Id: Texas 72
Collected 25 May 2011 in Medina county Texas, peridomestic/mixed Oak
Collected by Blake Sissel & Ed Wozniak, USA
Host: *Triatoma lecticularia*
MSH2

Position 381

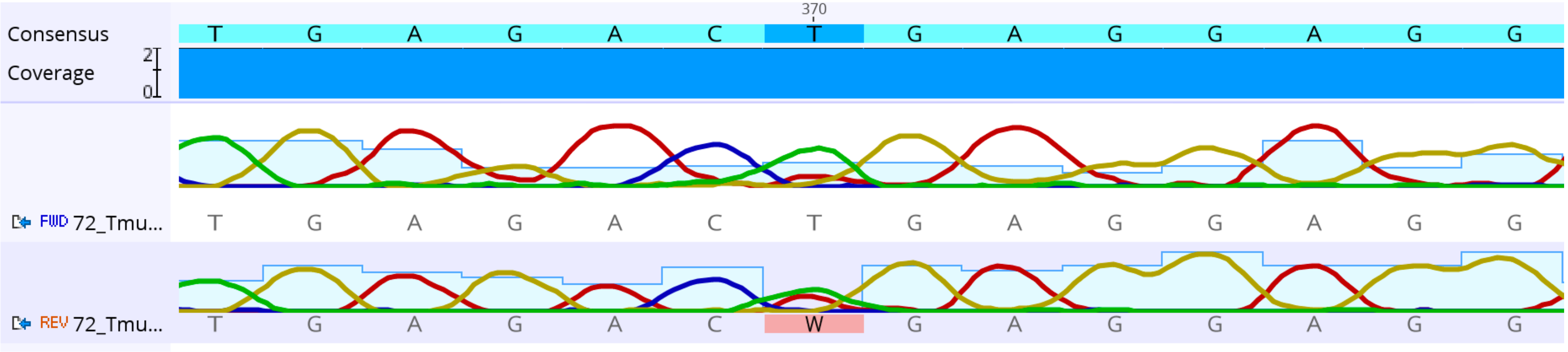


T: is from TcIV-USA allele
A: is from TcI allele

Potential Hybrid
Id: Texas 72
Collected 25 May 2011 in Medina county Texas, peridomestic/mixed Oak
Collected by Blake Sissel & Ed Wozniak, USA
Host: *Triatoma lecticularia*
MSH2

Position 399

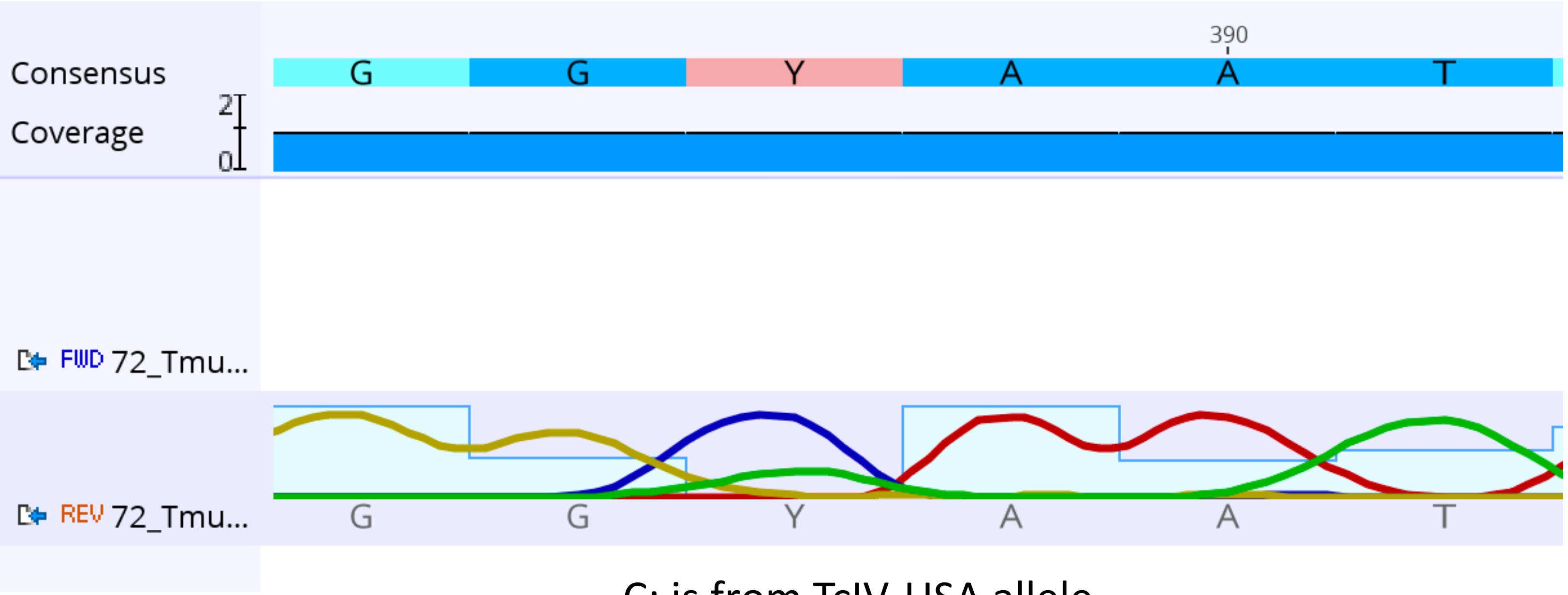


C: is from TcIV-USA allele
T: is from TcI allele

Potential Hybrid
Id: Texas 72
Collected 25 May 2011 in Medina county Texas, peridomestic/mixed Oak
Collected by Blake Sissel & Ed Wozniak, USA
Host: *Triatoma lecticularia*
MSH2

Position 421

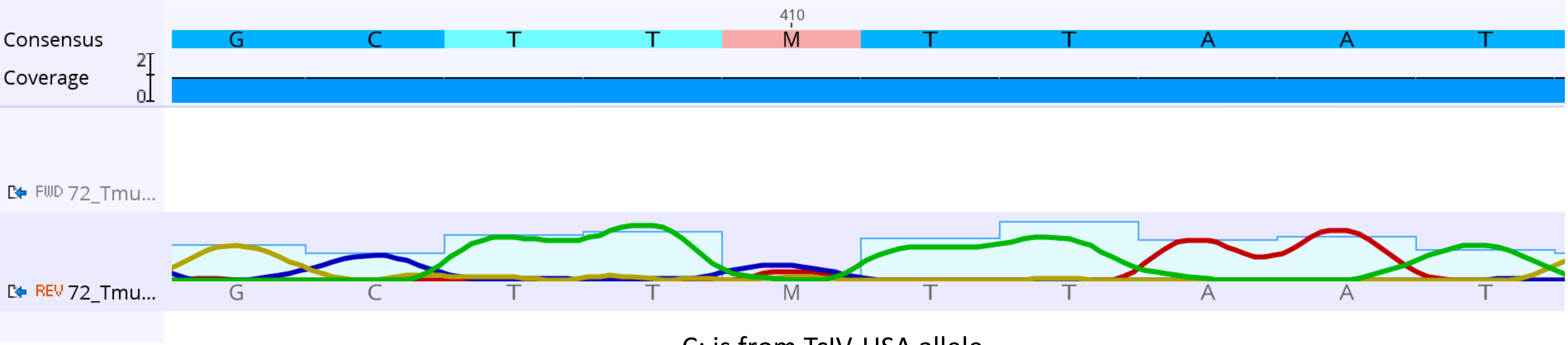


C: is from TcIV-USA allele
A: is from TcI allele

Potential Hybrid
Id: Texas 72
Collected 25 May 2011 in Medina county Texas, peridomestic/mixed Oak
Collected by Blake Sissel & Ed Wozniak, USA
Host: *Triatoma lecticularia*
MSH2

Position 555

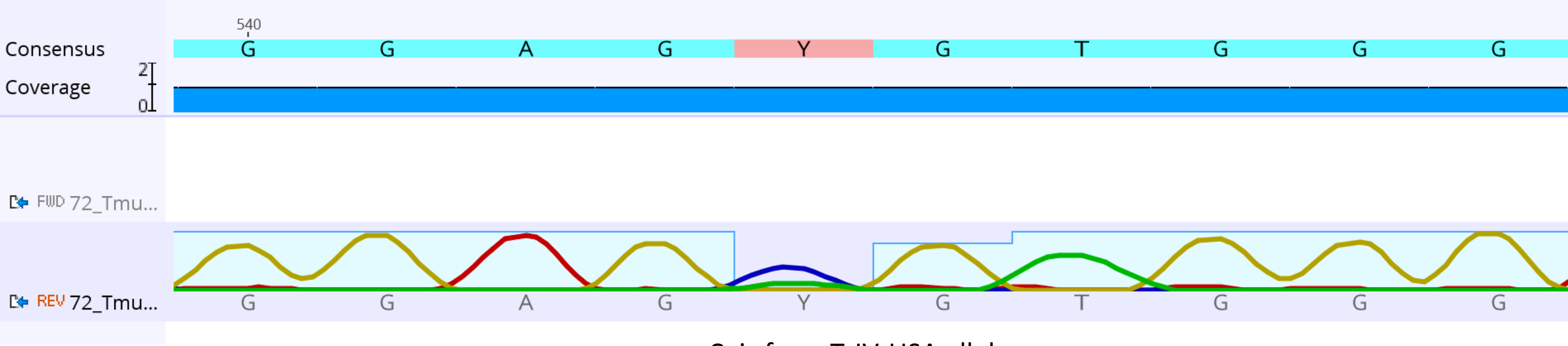


C: is from TcIV-USA allele
T: is from TcI allele

Potential Hybrid
Id: Texas 72
Collected 25 May 2011 in Medina county Texas, peridomestic/mixed Oak
Collected by Blake Sissel & Ed Wozniak, USA
Host: *Triatoma lecticularia*
MSH2

Position 566

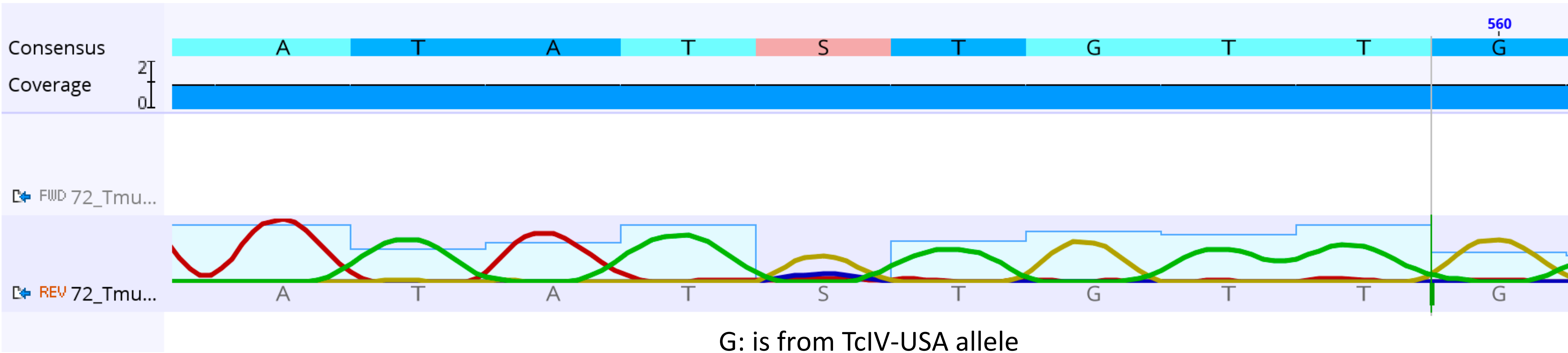


G: is from TcIV-USA allele
C: is from TcI allele
